# C-terminal truncation of CXCL10 attenuates inflammatory activity but retains angiostatic properties

**DOI:** 10.1101/2023.07.10.548382

**Authors:** Luna Dillemans, Karen Yu, Alexandra De Zutter, Sam Noppen, Mieke Gouwy, Nele Berghmans, Mirre De Bondt, Lotte Vanbrabant, Stef Brusselmans, Erik Martens, Dominique Schols, Pedro Elias Marques, Sofie Struyf, Paul Proost

**Affiliations:** Laboratory of Molecular Immunology, Department of Microbiology, Immunology and Transplantation, Rega Institute, KU Leuven, Leuven, Belgium; Laboratory of Virology and Chemotherapy, Department of Microbiology, Immunology and Transplantation, Rega Institute, KU Leuven, Leuven, Belgium; Laboratory of Immunobiology, Department of Microbiology, Immunology and Transplantation, Rega Institute, KU Leuven, Leuven, Belgium

**Author notes:** shared second author (Karen Yu, Alexandra De Zutter). CORRESPONDING AUTHOR Prof. dr. Paul Proost, Laboratory of Molecular Immunology, Department of Microbiology, Immunology and Transplantation, Rega Institute, KU Leuven. Address: Herestraat 49 box 1042, Leuven, Belgium.

**Keywords:** angiogenesis, chemokine, CXCL10, lymphocytes, posttranslational modifications, proteolysis, solid phase peptide synthesis

## Abstract

Interferon-γ-inducible protein of 10 kDa (IP-10/CXCL10) is a dual-function CXC chemokine that coordinates chemotaxis of activated T cells and natural killer (NK) cells via interaction with its G protein-coupled receptor (GPCR), CXC chemokine receptor 3 (CXCR3). As a consequence of natural posttranslational modifications, human CXCL10 exhibits a high degree of structural and functional heterogeneity. However, the biological effect of natural posttranslational processing of CXCL10 at the carboxy (C)-terminus has remained partially elusive. The truncated CXCL10 proteoform CXCL10_(1-73)_, lacking the four endmost C-terminal amino acids, was previously identified in human cell culture supernatant. To further explore the functioning of CXCL10_(1-73)_, we optimized its production in this study through Fmoc-based solid phase peptide synthesis (SPPS) and propose an SPPS strategy to efficiently generate human CXCL10 proteoforms. Compared to intact CXCL10_(1-77)_, CXCL10_(1-73)_ had diminished affinity for glycosaminoglycans including heparin, heparan sulfate and chondroitin sulfate A. Moreover, CXCL10_(1-73)_ exhibited an attenuated capacity to induce CXCR3A-mediated signaling, as evidenced in calcium mobilization assays and through quantification of phosphorylated extracellular signal-regulated kinase-1/2 (ERK1/2) and protein kinase B/Akt. Furthermore, CXCL10_(1-73)_ incited reduced primary human T lymphocyte chemotaxis *in vitro* and evoked less peritoneal ingress of CXCR3^+^ T lymphocytes in mice receiving intraperitoneal chemokine injections. In contrast, loss of the four endmost C-terminal residues did not affect the inhibitory properties of CXCL10 on spontaneous and/or FGF-2-induced migration, proliferation, wound healing, phosphorylation of ERK1/2, and sprouting of human microvascular endothelial cells. Thus, C-terminally truncated CXCL10 has attenuated inflammatory properties, but preserved anti-angiogenic capacity.

**GRAPHICAL ABSTRACT:** 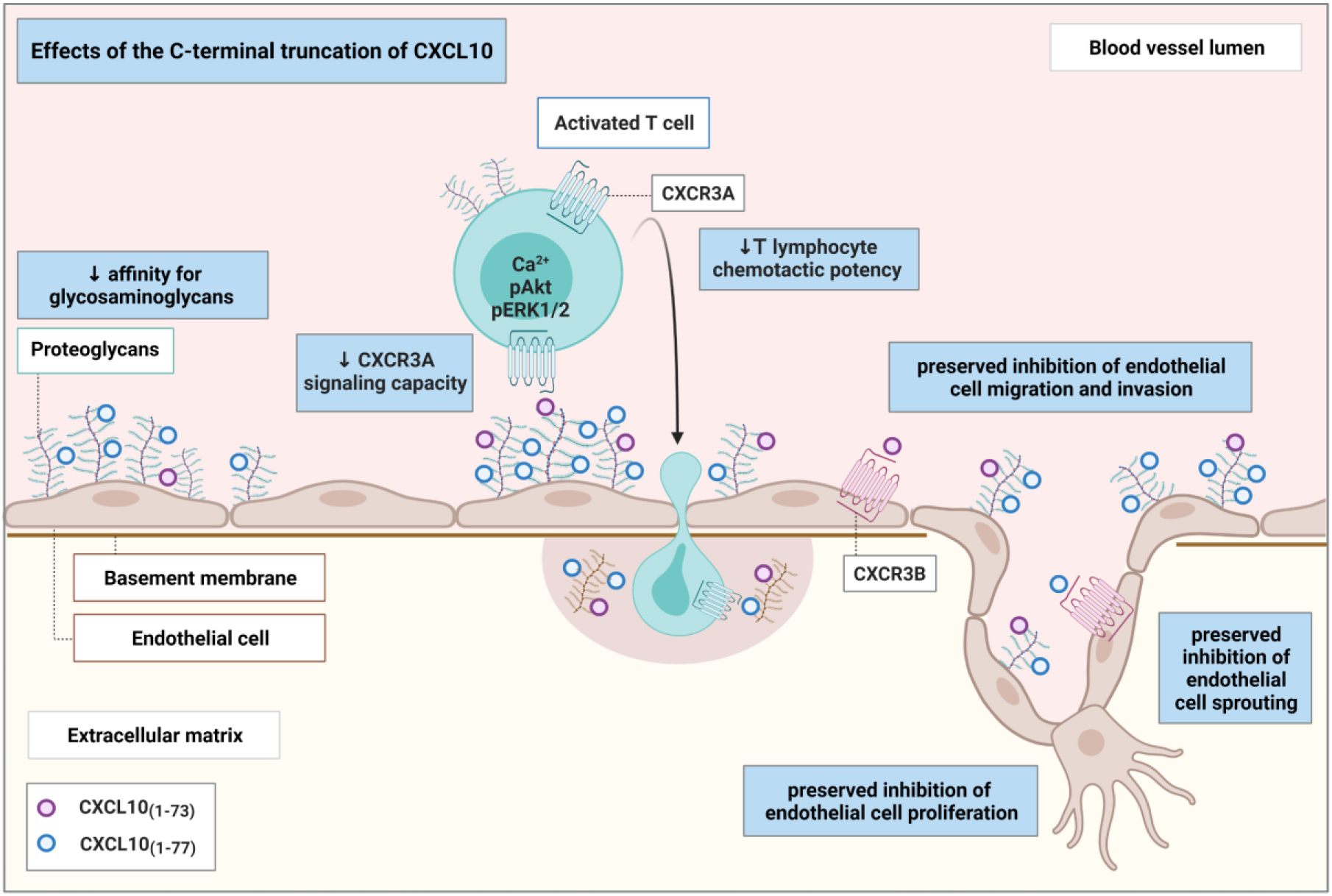

## INTRODUCTION

The superfamily of chemotactic cytokines or chemokines encompasses structurally similar, low molecular mass proteins (± 7-12 kDa) that govern directional leukocyte trafficking through interaction with chemokine-type G protein-coupled receptors (GPCRs) and glycosaminoglycans (GAGs) [1–4]. From a biochemical perspective, chemokines may be subdivided into four major subfamilies based on the number and positioning of the N-terminally located conserved cysteine residues [5–7]. CXC chemokines contain one random amino acid (‘X’) in between these cysteines and are further subcategorized based on the presence or absence of a Glu-Leu-Arg (‘ELR’) sequence located anterior of the CXC motif [6]. ELR^+^ CXC chemokines bind CXCR1 and/or CXCR2 and chemo-attract neutrophils, whereas ELR^−^ CXC chemokines primarily mediate directional lymphocyte trafficking [1,7,8]. Chemokines can also be categorized into functional subclasses, separating those with inflammatory as opposed to homeostatic actions, whereby ‘dual-function’ chemokines exhibit activities reminiscent of both inflammation and homeostasis [9,10]. Interferon-γ-inducible protein of 10 kDa (IP-10/CXCL10) is a dual-function ELR^−^ CXC chemokine that coordinates chemotaxis of activated CD4^+^ T_H_1 cells, CD8^+^ T cells, natural killer (NK) cells and NKT cells via interaction with its GPCR, CXC chemokine receptor 3 (CXCR3) [11–18]. In addition, CXCL10 exhibits potent anti-angiogenic activity [19–21].

Posttranslational modifications (PTMs) have been recognized as pivotal regulatory mechanisms that determine the chemokine functioning through modulating affinity and selectivity for GPCRs and GAGs [22–24]. PTMs are executed by specific enzymes that are co-expressed during inflammation [23]. These modifications include proteolytic truncation, glycosylation, nitration and citrullination, which may radically modify *in vitro* and *in vivo* chemokine potency [23]. CXCL10 is no exception to this rule and is highly susceptible to site-specific N- and C-terminal proteolytic processing [25,26]. In addition to intact CXCL10_(1-77),_ purification of natural CXCL10 from cell culture supernatant of stimulated human fibroblasts, primary keratinocytes, MG-63 osteosarcoma cells, human umbilical cord endothelial cells, and peripheral blood mononuclear cells (PBMC) revealed multiple natural CXCL10 proteoforms [27–33]. These natural isoforms of CXCL10 were missing four C-terminal amino acids (Lys^74^, Arg^75^, Ser^76^, and Pro^77^), lacking two to five N-terminal residues (Val^1^, Pro^2^, Leu^3^, Ser^4^, and Arg^5^) and/or containing a citrulline instead of Arg^5^ [27–31]. N-terminal truncation of CXCL10 by the enzyme dipeptidylpeptidase IV (DPPIV/CD26) generates CXCL10_(3-77)_, which functions as a chemotaxis antagonist with retained angiostatic properties [34]. The antagonistic activities of CXCL10_(3-77)_ in terms of lymphocyte chemotaxis were also demonstrated *in vivo* [35,36]. CD26 inhibition in C57BL/6 mice through sitagliptin administration restored lymphocyte-attracting activity of intraperitoneally injected human CXCL10, and of endogenous murine CXCL10 (mCXCL10), resulting in enhanced recruitment of CXCR3^+^ lymphocytes towards the peritoneal cavity and B16F10 melanoma tumors, respectively [35,36]. In addition, natural N-terminally truncated CXCL10_(3-77)_ was detected in plasma of patients with hepatitis C virus (HCV) infection and in urine of patients with bladder carcinoma and active tuberculosis [37–42]. Hence, these findings provide ample evidence that PTMs of CXCL10 have *in vivo* biological significance in experimental and clinical settings.

In contrast to N-terminal proteolysis, natural C-terminal truncation of human CXCL10 has been explored to a relatively limited extent, despite their verified presence in natural conditioned media of human cells [27,29,30]. The C-terminal residues have been previously implied in the anti-angiogenic and anti-parasitic properties of CXCL10 [43,44]. A C-terminal fragment of CXCL10, spanning the α-helical and coiled domain residues Pro^56^-Pro^77^, was reported to inhibit *in vitro* vascular endothelial growth factor (VEGF)-induced endothelial cell migration and *in vivo* vessel formation to a comparable extent as intact CXCL10_(1-77)_ [43]. Moreover, virulence factor glycoprotein-63 (GP63) of *Leishmania Major* cleaves off the C-terminal α-helix of CXCL10 at Ala^60^-Lys^62^, generating a CXCL10 proteoform with attenuated T cell chemotactic potential *in vitro* [44]. Interestingly, this potential immune evasion strategy may be shared with other intracellular pathogens including *Salmonella Typhimurium* and *Chlamydia trachomatis* [44]. In terms of structural modeling, C-terminal residues Ser^76^-Pro^77^ of human CXCL10 were not successfully modeled via NMR spectroscopy and crystallography [45,46]. Therefore, prediction of the precise location and proximity of the C-terminal amino acids relative to other residues in the peptide backbone of CXCL10 is still speculative, making structure-activity predictions challenging. C-terminal residues of mCXCL10 have been investigated to a more elaborate extent and were implicated in GAG and receptor binding [47–50].

The aim of the present study was to evaluate the effects of the naturally occurring C-terminal truncation of CXCL10 on the functional properties of this human chemokine. We introduced a strategy for Fmoc-based solid phase peptide synthesis (SPPS) of posttranslationally modified CXCL10 proteoforms with the example of CXCL10_(1-73)_. We utilized synthetic CXCL10_(1-73)_ to examine the effects of the C-terminal truncation. We discovered that the C-terminal truncation of CXCL10 attenuated the interaction with GAGs, the signaling properties through CXCR3A, and the ability to attract T lymphocytes *in vitro* and *in vivo*. However, the angiostatic properties of CXCL10, including the inhibition of migration, proliferation, wound healing, phosphorylation of extracellular signal-regulated kinase-1/2 (ERK1/2), and sprouting of endothelial cells, were not affected by the C-terminal processing.

## RESULTS

### Chemical synthesis of C-terminally truncated human CXCL10 proteoform CXCL10_(1-73)_

To investigate the biological properties of natural C-terminally truncated CXCL10_(1-73)_, we generated this chemokine synthetically. Initial SPPS was performed using standard reagents for chemokine synthesis [51,52]. When an Fmoc-Ser(But)-Wang resin was used with 2-(1H-7-Azabenzotriazol-1-yl)-1,1,3,3-tetramethyluronium hexafluorophosphate (HATU) and di-isopropylethylamine (DIEA) as a coupling system, a remarkably poor yield of correctly synthesized CXCL10_(1-73)_ (± 0.026%) was obtained (theoretical relative molecular mass [Mr] 8173.65, experimental Mr 8172.72) (Suppl. Fig. S1A). Since the aforementioned coupling reagents have been generally acknowledged to result in highly efficient coupling [53], we assumed that the synthesis failure may have been caused by the nature of the amino acid sequence of the protein itself. Hence, in the sequence of CXCL10_(1-73)_, 48% of the amino-acids were found to have hydrophobic side-chains. To prevent synthesis failure due to hydrophobicity, pseudoprolines were incorporated at key positions based on both theoretically predicted and experimentally determined challenging regions that required multiple deprotection steps during the failed synthesis [54]. Hence, pseudoproline dipeptides were inserted at Arg^5^-Thr^6^, Ile^12^-Ser^13^, Ala^43^-Thr^44^ and Val^68^-Ser^69^ in the peptide backbone of CXCL10. In addition, 1,1,3,3-tetramethyluronium hexafluorophosphate (HCTU) and 4-methylmorpholine (NMM) were used as alternative high quality coupling reagents [55–57]. Despite these modifications in the SPPS, a very low amount of successfully generated CXCL10_(1-73)_ was present upon analysis of the synthetic material. One major and highly abundant contaminant was detected, i.e. N-terminally shortened acetylated CXCL10_(31-73)_ (theoretical Mr 4870.78, experimental Mr 4868.86) (**Fig. S1B**). The acetylation clearly points towards a synthesis artefact, resulting from impaired peptide amide bond formation between Ile^30^ and Pro^31^. Coupling of amino acids to C-terminally resin-attached Pro residues is often more challenging given the reduced reactivity of the secondary amine located in the proline oxazolidine ring structure. The formation of this shortened peptide was circumvented via the selective incorporation of a specific dipeptide building block at Ile^30^-Pro^31^, i.e. Fmoc-L-Ile-L-Pro (**Fig. S1C**). Subsequently, the successfully obtained purified crude linear material was folded, as CXCL10 contains two disulfide bridges (Cys^9^-Cys^36^ and Cys^11^-Cys^53^). Initial folding was performed through incubation in 150 nM tris(hydroxymethyl)-aminomethane (Tris; pH 8.6) supplemented with 3 mM ethylenediaminetetraacetic acid (EDTA), 0.3 mM reduced glutathione (GSH) 3 mM oxidized glutathione (GSSG), and 1 M guanidine hydrochloride for 5 h under continuous rotation [51]. However, since two glutathione residues (each with Mr 307.33) remained covalently coupled to the cysteines in the peptide backbone after the folding procedure (theoretical Mr 8788.31, experimental Mr 8785.02) (**Fig. S1D**), this approach resulted in a significant portion of the synthetic protein being incompletely folded. Therefore, an alternative folding methodology was applied in which the crude linear protein was incubated in 1.0 M guanidine hydrochloride and 0.1 M Tris (pH 8.5) whilst continuously stirred in air for 24 h to allow formation of the disulfide bridges [58]. Using this strategy of concomitant use of a hydrophilic resin, pseudoproline dipeptides, a dipeptide building block at pivotal positions, high quality hydrophobic solvents and oxidative folding under exposure of air (**Fig. 1**), correctly synthesized and folded CXCL10_(1-73)_ was obtained (**Fig. 2**; theoretical Mr 8173.65, experimental Mr 8172.37).

**FIGURE 1.**
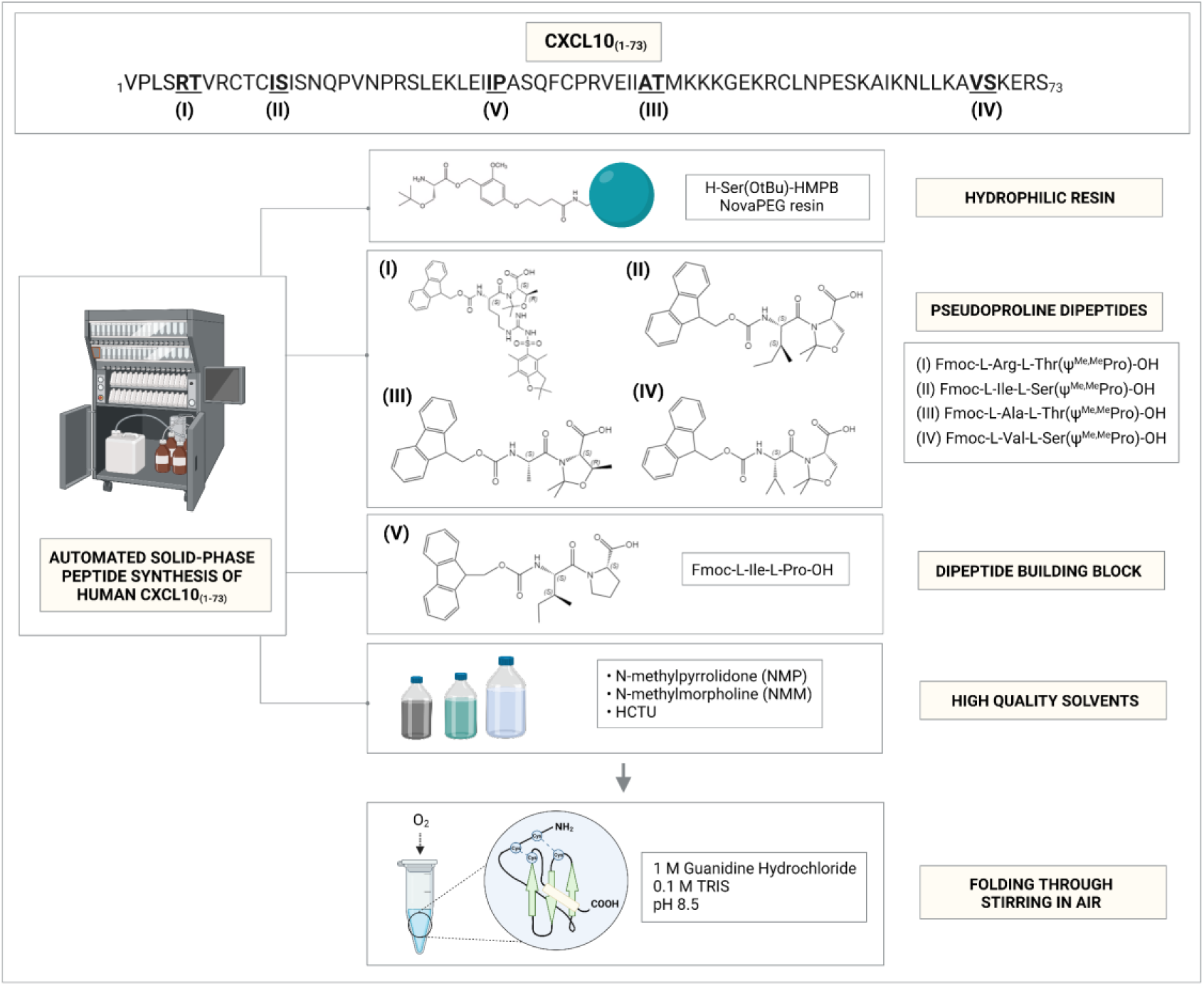
Strategy for successful solid phase peptide synthesis (SPPS) of human C-terminally truncated CXCL10_(1-73)_ via Fmoc chemistry. Four crucial aspects were defined to ensure a successful SPPS using Fmoc chemistry including the combined use of a hydrophilic resin, pseudoproline dipeptides and a dipeptide building block (indicated I-IV) at crucial positions and the continuous application of high quality solvents. The peptide backbone of CXCL10_(1-73)_ is depicted with the key positions at which pseudoproline dipeptides and the dipeptide building block (bold and underlined) were incorporated. Figure was created with Biorender. Fmoc, 9-fluorenylmethoxycarbonyl; HCTU, O-(1H-6-chloro-benzotriazole-1-yl)-1,1,3,3-tetramethyluronium hexafluorophosphate; HMPB, 4-(4-hydroxymethyl-3-methoxyphenoxy)butyric acid; NMM, 4-methylmorpholine; NMP, N-methyl-2-pyrrolidone; OtBu, tert-butyl ester; PEG, polyethylene glycol.

**FIGURE 2.**
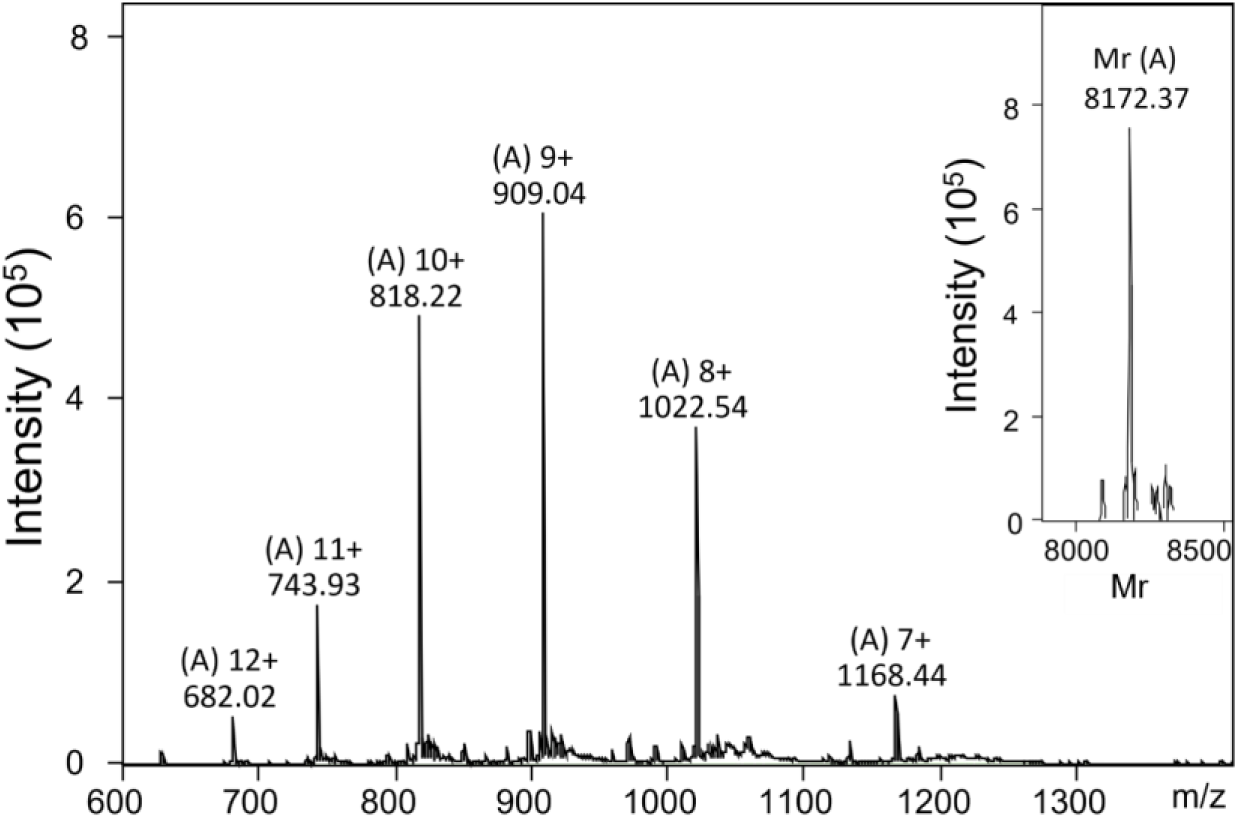
Averaged mass spectrum of pooled and folded synthetic CXCL10_(1-73)_. The intensity of the detected ions with their respective mass/charge (m/z) ratio are displayed. The relative molecular mass (Mr) of CXCL10_(1-73)_ was calculated with Bruker deconvolution software (inset on the right) based on the detected ions with their respective m/z ratio, i.e. the ions marked by [A] with 7 to 12 positive charges. This experimental Mr (8172.37) corresponded to the calculated theoretical Mr (8173.65).

### CXCL10_(1-73)_ has reduced affinity for GAGs compared to native CXCL10_(1-77)_

Given the potential involvement of the positively charged C-terminal amino-acids Lys^74^ and Arg^75^ of CXCL10 in binding to GAGs, we investigated binding of CXCL10_(1-73)_ to heparin, heparan sulfate (HS) and chondroitin sulfate (CS)-A using surface plasmon resonance (SPR) technology. CXCL4 was included as a positive control for CS-A binding [59]. Varying concentrations of CXCL10_(1-77)_, CXCL10_(1-73)_ and CXCL4 were sent over a neutravidin-coated CM4 chip on which distinct GAGs were immobilized in individual flow channels (**Fig. 3A-H**). To characterize the nature of the interaction, kinetic parameters were determined from the association (1 to 120 seconds) and dissociation (120 to 300 seconds) phases of the SPR sensorgrams (**Table 1**). Binding kinetics were analyzed and fitted through the 1:1 binding model with mass transfer correction [60,61] to calculate apparent K_D_ values (**Fig. S2**). CXCL10_(1-73)_ exhibited 3.7-fold decreased affinity for HS compared to CXCL10_(1-77)_ (**Fig. 3A and 3D, Table 1**). CXCL10_(1-73)_ showed 32.4-fold reduced affinity for heparin compared to CXCL10_(1-77)_ (**Fig. 3C and 3F, Table 1**). Furthermore, CXCL10_(1-73)_ bound weakly and 15.3-fold less efficient to CS-A compared to CXCL10_(1-77)_ (**Fig. 3B and 3E, Table 1**). As expected, CXCL4 exhibited higher affinity for CS-A compared to CXCL10_(1-77)_ (**Fig. 3H, Table 1**). Thus, CXCL10_(1-73)_ displayed diminished affinity for HS, CS-A and heparin compared to CXCL10_(1-77)_.

**FIGURE 3.**
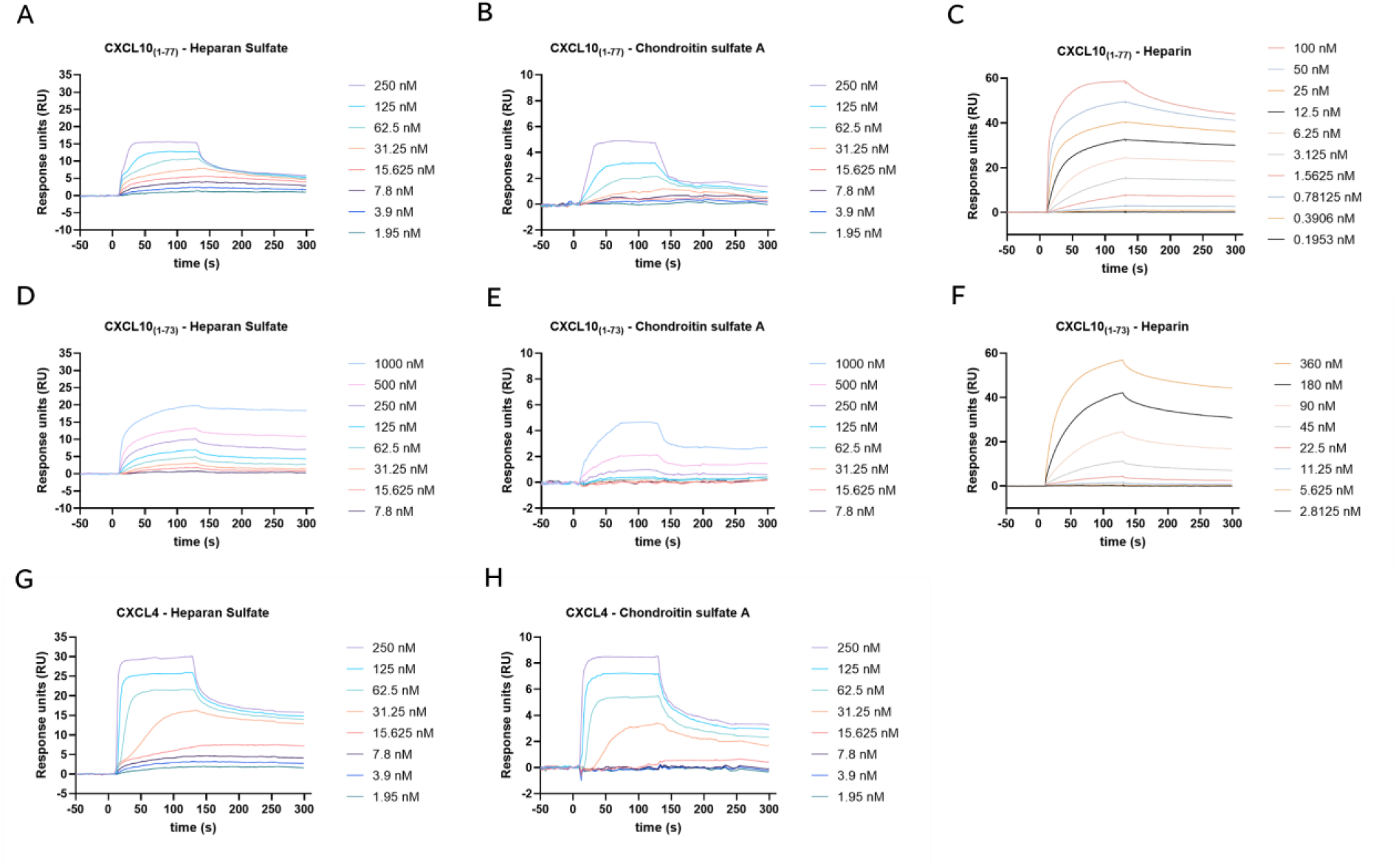
Glycosaminoglycan affinity of C-terminally truncated CXCL10_(1-73)_ is reduced compared to intact CXCL10_(1-77)_. CXCL10_(1-77)_ and CXCL10_(1-73)_ at varying concentrations were sent over the neutravidin-coated CM4 Biosensor chip surfaces on which biotinylated heparin, HS or CS-A were immobilized. Representative SPR sensorgrams are shown (from 4 independent experiments) displaying the affinity for heparin, HS and CS-A of (**A-C**) CXCL10_(1-77)_, (**D-F**) CXCL10_(1-73)_ and (**G-H**) CXCL4. SPR sensorgrams were obtained after subtracting the baseline signal of the reference channel and a blank of the respective channel. The following concentrations were used in ½ serial dilution series for curve fitting: CXCL10_(1-77)_ at 100 nM to 0.195 nM for heparin and 250 nM to 1.953 nM for HS and CS-A; CXCL10_(1-73)_ at 360 nM to 2.813 nM for heparin and 1000 nM to 7.813 nM for HS and CS-A; CXCL4 at 250 nM to 1.953 nM for HS and CS-A. Kinetic parameters were determined from the association phase (1 to 120 seconds) and dissociation phase (120 to 300 seconds) of the SPR sensorgrams. The y-axis displays the SPR response in response units (RU).

**TABLE 1.** Kinetic parameters of the interaction between human CXCL10 proteoforms and GAGs

| Chemokine | Heparan sulfate |  |  | Heparin |  |  | Chondroitin sulfate A |  |  |
| --- | --- | --- | --- | --- | --- | --- | --- | --- | --- |
| | $k_{on}$ (1/M.s) | $k_{off}$ (1/s) | Apparent $K_D$<br>(nM) | $k_{on}$ (1/M.s) | $k_{off}$ (1/s) | Apparent $K_D$<br>(nM) | $k_{on}$ (1/M.s) | $k_{off}$ (1/s) | Apparent $K_D$<br>(nM) |
| **CXCL4** | (8.58 ± 0.54)<br>E+05 | (3.21 ± 0.14)<br>E-03 | 3.75 ± 0.10 | N.D. | N.D. | N.D. | (4.36 ± 0.60)<br>E+05 | (6.71 ± 0.77)<br>E-03 | 15.91 ± 2.61 |
| **CXCL10<sub>(1-77)</sub>** | (5.28 ± 0.84)<br>E+05 | (3.41 ± 0.43)<br>E-03 | 6.73 ± 0.65 | (9.76 ± 0.41)<br>E+05 | (1.00 ± 0.04)<br>E-03 | 1.03 ± 0.01 | (1.69 ± 0.09)<br>E+05 | (5.57 ± 0.34)<br>E-03 | 33.40 ± 2.86 |
| **CXCL10<sub>(1-73)</sub>** | (4.86 ± 0.96)<br>E+04 | (1.26 ± 0.27)<br>E-03 | 25.23 ± 1.16 | (4.26 ± 0.32)<br>E+04 | (1.39 ± 0.04)<br>E-03 | 33.42 ± 2.65 | (3.15 ± 1.08)<br>E+03 | (1.06 ± 0.12)<br>E-03 | 512.40 ± 191.31 |

Values represent the mean ± SEM of 3 to 4 independent experiments. Kinetic parameters were determined from the association phase (1 to 120 seconds) and dissociation phase (120 to 300 seconds) of the SPR sensorgrams. The $K_D$ was calculated from the ratio of $k_{off}$ over $k_{on}$ (nM) determined by the 1:1 binding model with mass transfer correction. $k_{on}$ association rate constant ( $M^{-1} s^{-1}$ ); $k_{off}$ dissociation rate constant ( $s^{-1}$ ); $K_D$ dissociation equilibrium (affinity) constant. N.D., not determined.

### CXCL10_(1-73)_ is a less potent inducer of second messenger signaling downstream of CXCR3A compared to native CXCL10_(1-77)_

In calcium assays, CXCR3A-transfected CHO cells were used to determine whether CXCL10_(1-73)_ had similar potency as CXCL10_(1-77)_ to induce mobilization of intracellular calcium. Only highly elevated concentrations of CXCL10_(1-73)_ (i.e., 135 nM and 270 nM) were able to induce a moderate increase in intracellular calcium concentrations, thereby reaching comparable calcium levels as upon stimulation with 1 nM CXCL10_(1-77)_ (**Fig. 4A**). At 3 nM of CXCL10_(1-77)_, calcium mobilization was even significantly higher compared to 270 nM of CXCL10_(1-73)_. We also observed that the time between administration of the stimulus and the initiation of the calcium increase was prolonged for CXCL10_(1-73)_ independent of the administrated dose (**Fig. 4B**). The markedly limited capacity of CXCL10_(1-73)_ compared to CXCL10_(1-77)_ to mobilize intracellular calcium sparks the notion of potential CXCR3 desensitization by this C-terminally shortened CXCL10 proteoform at inactive concentrations. Indeed, when 100 seconds prior to a stimulus of 3 nM CXCL10_(1-77)_, 45 nM or 90 nM of inactive CXCL10_(1-73)_ was added to the cells, the increase of intracellular calcium upon stimulation with CXCL10_(1-77)_ was reduced by 69.9% and 72.2%, respectively (**Fig. 4C-D**). CXCR3 desensitization may be due to partial agonism or receptor internalization. Therefore, we evaluated the effects of both CXCL10 proteoforms on CXCR3 internalization on primary T lymphocytes derived from PBMCs of individual donors and stimulated with phytohemagglutinin (PHA)- and IL-2. CXCR3 expression was corroborated on these primary T lymphocytes (**Fig. S3**). Most T lymphocytes (median expression 68.6%, median fluorescence intensity [MFI] 1877) were positive for CXCR3 (**Fig. 4E**). CXCL10_(1-77)_ induced CXCR3 internalization more potently compared to CXCL10_(1-73),_ as the relative surface expression of CXCR3A was significantly further reduced upon incubation with 30 nM CXCL10_(1-77)_ compared to 45 nM CXCL10_(1-73)_. In addition, we found that relatively limited internalization of CXCR3 was induced by concentrations of CXCL10_(1-73)_ that were able to desensitize CXCR3A, i.e. 45 nM (mean MFI CXCR3 expression 90.1%) and 90 nM (mean MFI CXCR3 expression 83.3%) (**Fig. 4F**). Hence, receptor internalization and partial agonism likely both contribute to CXCR3A desensitization mediated by CXCL10_(1-73)_.

**FIGURE 4.**
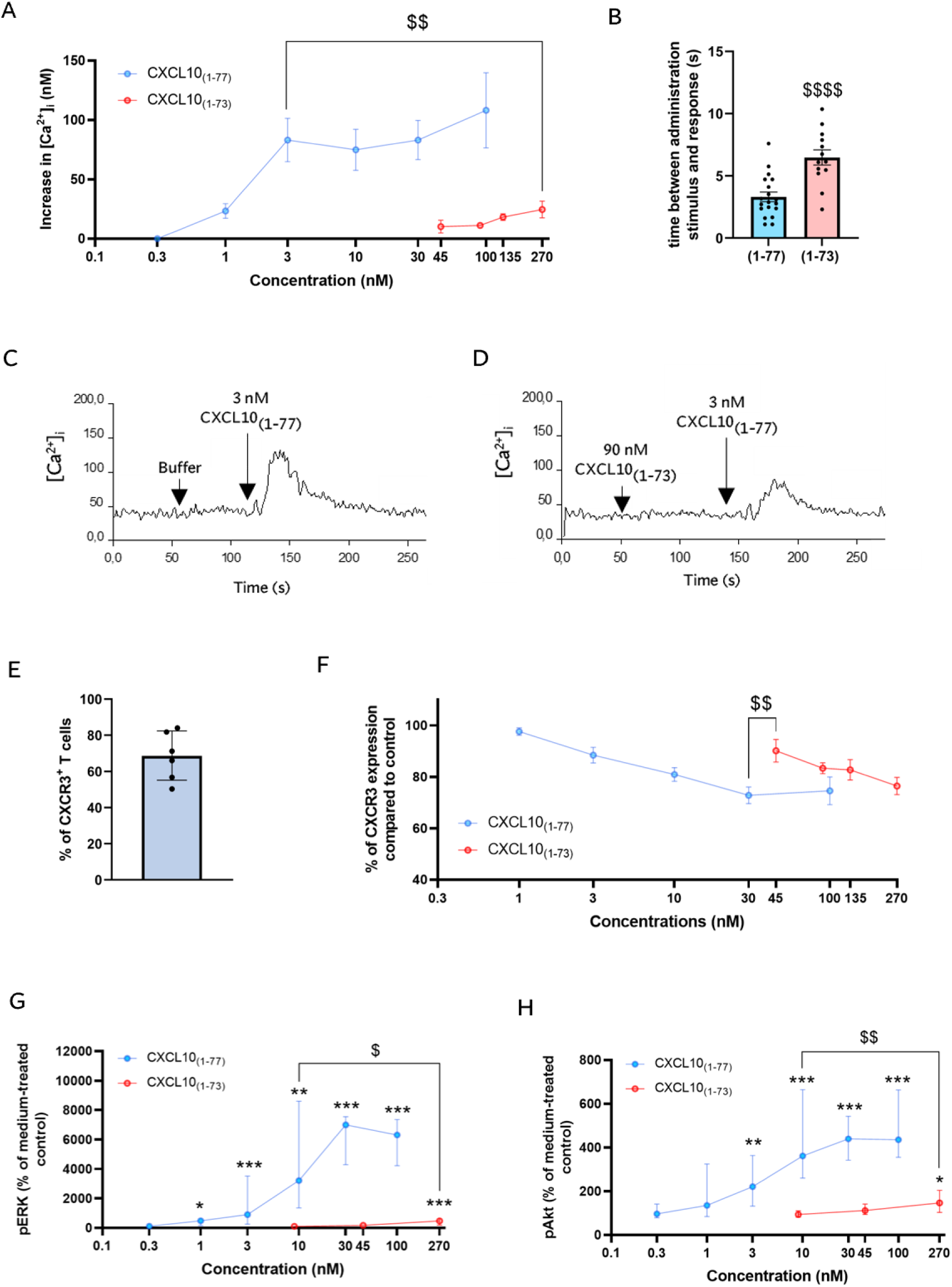
C-terminally truncated CXCL10_(1-73)_ has less potent signaling capacity in CXCR3A-transfected CHO cells and PHA- and IL-2-stimulated T lymphocytes compared to intact CXCL10_(1-77)_. (**A**) CXCL10_(1-77)_ (0.3, 1, 3, 10, 30 and 100 nM) and CXCL10_(1-73)_ (45, 90, 135 and 270 nM) were evaluated for their ability to induce an increase of the intracellular calcium concentration (indicated as [Ca^2+^]_i_) in CXCR3A-transfected CHO cells. Results are displayed as mean (± SEM) increase of the intracellular calcium concentration of 4 independent experiments. Responses induced by 3 nM CXCL10_(1-77)_ and 270 nM CXCL10_(1-73)_ were compared using an unpaired t-test ($$ p < 0.01). (**B**) Time between administration of the stimulus and response in sec (s). Results are displayed as mean (± SEM) of 3 independent experiments with 19 measurements for CXCL10_(1-77)_ and 13 measurements for CXCL10_(1-73)_. Statistically significant differences between CXCL10_(1-77)_ and CXCL10_(1-73)_ were determined by an unpaired t-test ($$$$ p < 0.0001). (**C-D**) Representative curves show desensitization of CXCR3A-mediated intracellular calcium mobilization upon stimulation with 3 nM CXCL10_(1-77)_ following treatment with CXCL10_(1-73)_ or buffer as first stimulus. (**E**) CXCR3 expression on PHA- and IL-2-stimulated T lymphocytes (gated as CD3^+^ CD56^−^ CD19^−^ cells) was evaluated through flow cytometry with proportions of CXCR3^+^ T lymphocytes. (**F**) Relative surface expression of CXCR3 on primary T lymphocytes (compared to medium-treated control cells) following stimulation with CXCL10_(1-77)_ (1, 3, 10, 30 and 100 nM) and CXCL10_(1-73)_ (45, 90, 135 and 270 nM). Results are shown as median (± SEM) of 3 independent experiments with 9 different cell preparations in total. Responses induced by 30 nM CXCL10_(1-77)_ and 45 nM CXCL10_(1-73)_ were compared using an unpaired t-test ($$ p < 0.01). (**G-H**) The levels of ERK1/2 and Akt phosphorylation in CXCR3A-transfected CHO cells stimulated with CXCL10 proteoforms were determined by DuoSet ELISAs. Cells were stimulated with different concentrations of CXCL10_(1-77)_ (0.3, 1, 3, 10, 30 and 100 nM) or CXCL10_(1-73)_ (9, 45 and 270 nM). Results are shown as median (± IQR) of 4 to 8 independent experiments. Statistically significant ERK1/2 and Akt phosphorylation induced by CXCL10_(1-77)_ and CXCL10_(1-73)_ compared to medium-treated cells were determined by Mann-Whitney U test (* p < 0.05, **, p <0.01, *** p < 0.001). Comparison of the ERK1/2 and Akt phosphorylation induced by 10 nM CXCL10_(1-77)_ and 270 nM CXCL10_(1-73)_ was also performed through a Mann-Whitney U test ($ p < 0.05, $$, p < 0.01).

Second, the ability of CXCL10_(1-73)_ to induce phosphorylation of ERK1/2 and protein kinase B/Akt in CXCR3A-transfected CHO cells was tested. Similar to the calcium signaling experiments, high concentrations of CXCL10_(1-73)_ (270 nM) only weakly induced phosphorylation of ERK1/2 and Akt, thereby inciting comparable median levels of phosphorylated ERK1/2 (pERK1/2) and phosphorylated Akt (pAkt) as induced by 1 nM of CXCL10_(1-77)_ (**Fig. 4G-H**). Furthermore, significantly higher pAkt and pERK levels were observed upon treatment with 10 nM of CXCL10_(1-77)_ compared to 270 nM of CXCL10_(1-73)_ (**Fig. 4G-H**). Hence, CXCL10_(1-73)_ exhibited reduced potency compared to CXCL10_(1-77)_ to induce intracellular calcium mobilization, internalization of CXCR3A and phosphorylation of ERK1/2 and Akt.

### CXCL10_(1-73)_ evokes less chemotactic migration of primary lymphocytes compared to native CXCL10_(1-77)_

Given that the C-terminal truncation of CXCL10 significantly attenuated CXCR3 signaling, we further investigated whether CXCL10_(1-73)_ also exhibited reduced T lymphocyte chemotactic capacities. To ascertain adequate responsiveness of T lymphocytes, CXCL12α was included as positive control. PHA- and IL-2-stimulated T lymphocytes are known to express CXCR4 and pronouncedly migrate after exposure to CXCL12α [62]. Most T lymphocytes (median expression 81.6%, median MFI 2303) were positive for CXCR3 (**Fig. 5A-B**). Given that *in vitro* T cell chemotaxis induced by CXCL10 has been shown to occur in the absence of coating [47] and to exclude that distinct binding affinities of the CXCL10 proteoforms to extracellular matrix proteins underly the difference in T lymphocyte migration, we evaluated migration through uncoated membranes. We confirmed that chemotaxis was significantly and dose-dependently increased upon stimulation with CXCL12α relative to spontaneous migration of medium-treated cells (**Fig. 5C**). Starting from 1 nM, CXCL10_(1-77)_ induced a significant and dose-dependent migration of CXCR3^+^ T lymphocytes compared to cells exposed to buffer. The migratory response of T lymphocytes towards CXCL10_(1-73)_ was significantly increased compared to buffer only from 10 nM CXCL10_(1-73)_ onwards. Chemotaxis of T cells was significantly diminished for CXCL10_(1-73)_ compared to CXCL10_(1-77)_ at all tested concentrations (i.e. 1 nM, 3 nM, 10 nM, 30 nM and 100 nM).

**FIGURE 5.**
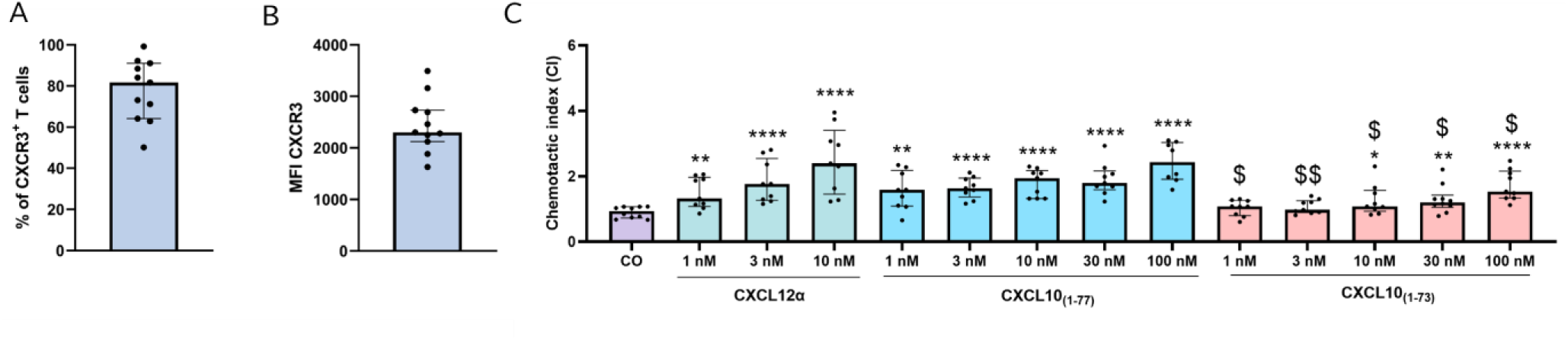
Migration of PHA- and IL-2-stimulated CXCR3^+^ T lymphocytes towards C-terminally truncated CXCL10_(1-73)_ is reduced compared to CXCL10_(1-77)_. CXCR3 expression on PHA- and IL-2-stimulated T lymphocytes (gated as CD3^+^ CD56^−^ CD19^−^ cells) was evaluated through flow cytometry with (**A**) proportions of CXCR3^+^ T lymphocytes and (**B**) MFI of CXCR3. (**C**) Chemotactic index (CI) showing migration of PHA- and IL-2-stimulated T lymphocytes through membranes without coating after treatment with medium (HBSS + 0.1% BSA) as control condition (CO) or serial dilution of CXCL10_(1-77)_ (100 nM to 1 nM) or CXCL10_(1-73)_ (100 nM to 1 nM). Results are shown as median (± IQR) of 4 independent experiments with 10 different cell preparations in total. Statistically significant chemotactic indices compared to medium-treated cells (* p < 0.05, **, p <0.01, *** p < 0.001, **** p < 0.0001) and between CXCL10 proteoforms ($ p < 0.05, $$ p < 0.01) were determined by Mann-Whitney U test

CXCL10 requires presentation on GAGs to mediate transendothelial migration of primary human CD4^+^ T lymphocytes under conditions of physiological shear stress [63]. Hence, we also evaluated migration through membranes coated with different extracellular matrix proteins (i.e. bovine fibronectin, human FN and human type I collagen). Using these different proteoglycans, CXCL10_(1-73)_ also induced significantly less T lymphocyte migration compared to CXCL10_(1-77)_ at 10 nM, 30 nM and/or 100 nM (**Fig. S4A-C**) with no significant differences between human FN and human type I collagen. Thus, in line with the observation of the signaling assays, C-terminal processing of CXCL10 also significantly attenuates its chemotactic properties on primary CXCR3^+^ T lymphocytes.

### CXCL10_(1-73)_ is equally potent in exerting antiangiogenic actions compared to native CXCL10_(1-77)_

Since a CXCL10-derived peptide CXCL10_(56-77)_ was equally potent in mediating angiostatic effects as intact CXCL10_(1-77)_ [43], one could presume that CXCL10_(1-73)_ has attenuated capacity to exert anti-angiogenic actions. For this reason, we examined the activities of CXCL10_(1-73)_ on human microvascular endothelial cells (HMVEC) in migration, proliferation, wound healing, signal transduction and sprouting assays.

First, chemotactic migration of endothelial cells in the presence of CXCL10 proteoforms was evaluated. Migration of HMVEC from an upper towards the lower chamber was monitored and analyzed at 12 h to exclude potential anti-proliferative effects of CXCL10. Stimulation with FGF-2 caused a significant increase of migration of endothelial cells to the lower chamber (**Fig. 6A**). CXCL10_(1-73)_ was equally potent as intact CXCL10_(1-77)_ in inhibiting FGF-2-induced chemotaxis of HMVEC. Starting from 12 nM, both CXCL10 isoforms suppressed FGF-2-mediated migration of endothelial cells in a dose-dependent manner. Accordingly, spontaneous HMVEC chemotaxis was dose-dependently inhibited with similar efficiency for both CXCL10 proteoforms from a concentration of 120 nM onwards (**Fig. 6B**). Although spontaneous migration was slightly decreased at 12 nM of CXCL10_(1-77)_, both CXCL10 proteoforms were not able to significantly counteract spontaneous chemotaxis of endothelial cells at a dose of 12 nM in contrast to the FGF-2-induced migration.

**FIGURE 6.**
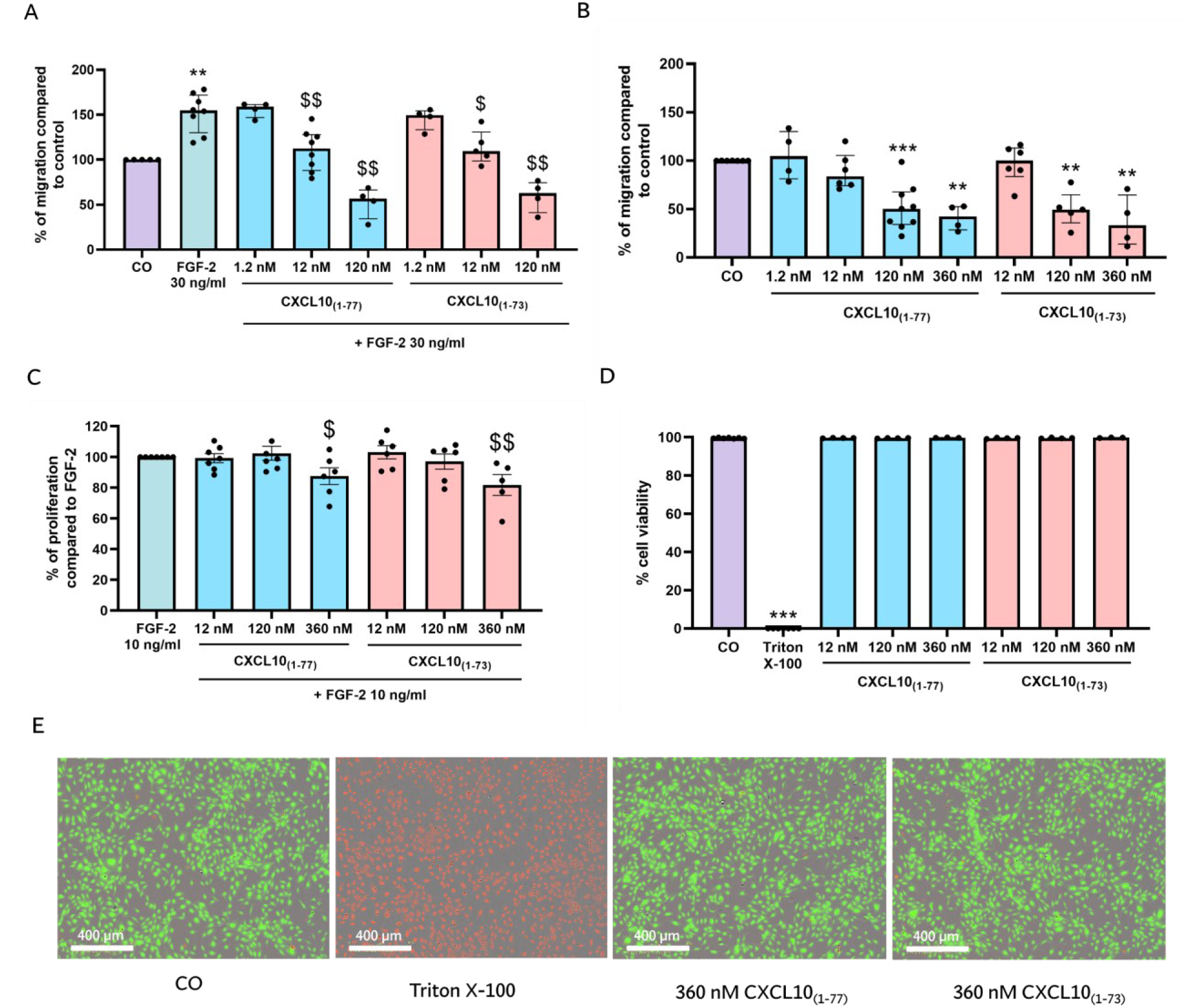
Equipotent inhibition of spontaneous and FGF-2-induced HMVEC migration by intact CXCL10_(1-77)_ or C-terminally truncated CXCL10_(1-73)_ without exerting cellular toxicity. HMVEC chemotaxis was measured towards (**A**) 30 ng/ml FGF-2 and (**B**) EBM-2 + 0.4% FCS treated cells (CO) in the presence or absence of CXCL10_(1-77)_ or CXCL10_(1-73)_. The data are displayed as median (± IQR) of 4 to 7 independent experiments. Statistically significant differences in migration compared to cells treated with control medium or FGF-2 were determined by a Mann-Whitney U test (** p < 0.01, *** p < 0.001 for comparison to control, $$ p < 0.01 for comparison to FGF-2). (**C**) FGF-2-induced proliferation of HMVEC was examined in the presence or absence of CXCL10_(1-77)_ or CXCL10_(1-73)_. The data are displayed as mean (± SEM) of 5 to 7 independent experiments. Statistically significant differences in proliferation compared to cells treated with 10 ng/ml FGF-2 were determined by an unpaired t-test ($ p < 0.05; $$ p < 0.01). (**D**) Cellular toxicity was assessed after 30 h of stimulation with CXCL10_(1-77)_ or CXCL10_(1-73)_. The median (± IQR) of 3 to 4 independent experiments is shown. Statistically significant differences in cell viability compared to cells treated with control were determined by a Mann-Whitney U test (*** p < 0.001). (**E**) Representative images are displayed of HMVEC treated with control medium (CO; MCDB131 + 0.4% [*v/v*] FCS), 2% (*v/v*) Triton X-100 to induce cell death, and 360 nM of CXCL10_(1-77)_ or CXCL10_(1-73)_. Scale bar = 400 µm.

Second, we examined the effects of both CXCL10 proteoforms on proliferation of HMVEC. Both CXCL10_(1-73)_ and CXCL10_(1-77)_ equally inhibited FGF-2-induced proliferation at 360 nM (**Fig. 6C**). To ascertain that the inhibitory effects of CXCL10 proteoforms were not due to cellular toxicity, their effects on endothelial cell survival were investigated. At the highest evaluated concentration (360 nM), CXCL10_(1-73)_ and CXCL10_(1-77)_ did not affect the viability of HMVEC after incubation for 30 h (**Fig. 6D-E**).

Third, we assessed the influence of CXCL10_(1-73)_ and CXCL10_(1-77)_ on the ability of endothelial cells to spontaneously invade and migrate into a scratch wound in a confluent monolayer. At a dose of 120 nM, CXCL10_(1-73)_ and CXCL10_(1-77)_ suppressed spontaneous wound healing, marked by significantly attenuated relative wound density and wound confluence (**Fig. 7A-B**). Both CXCL10 proteoforms were not able to significantly suppress spontaneous migration and invasion of endothelial cells at a dose of 12 nM. Differences in wound confluence were also observed after imaging the wound area (**Fig. 7C-D; Fig. S5**). In addition, CXCL10 proteoforms suppressed FGF-2-induced wound healing at 360 nM, but not at 120 nM (**Fig. S6A-C**). The apparent difference of the effects of CXCL10 proteoforms on FGF-2-induced migration at 12 nM observed through the xCELLigence migration assay may be explained by the limited resolution of the wound healing assay as opposed to the migration assay (i.e. accurate measurement of electrical impedance).

**FIGURE 7.**
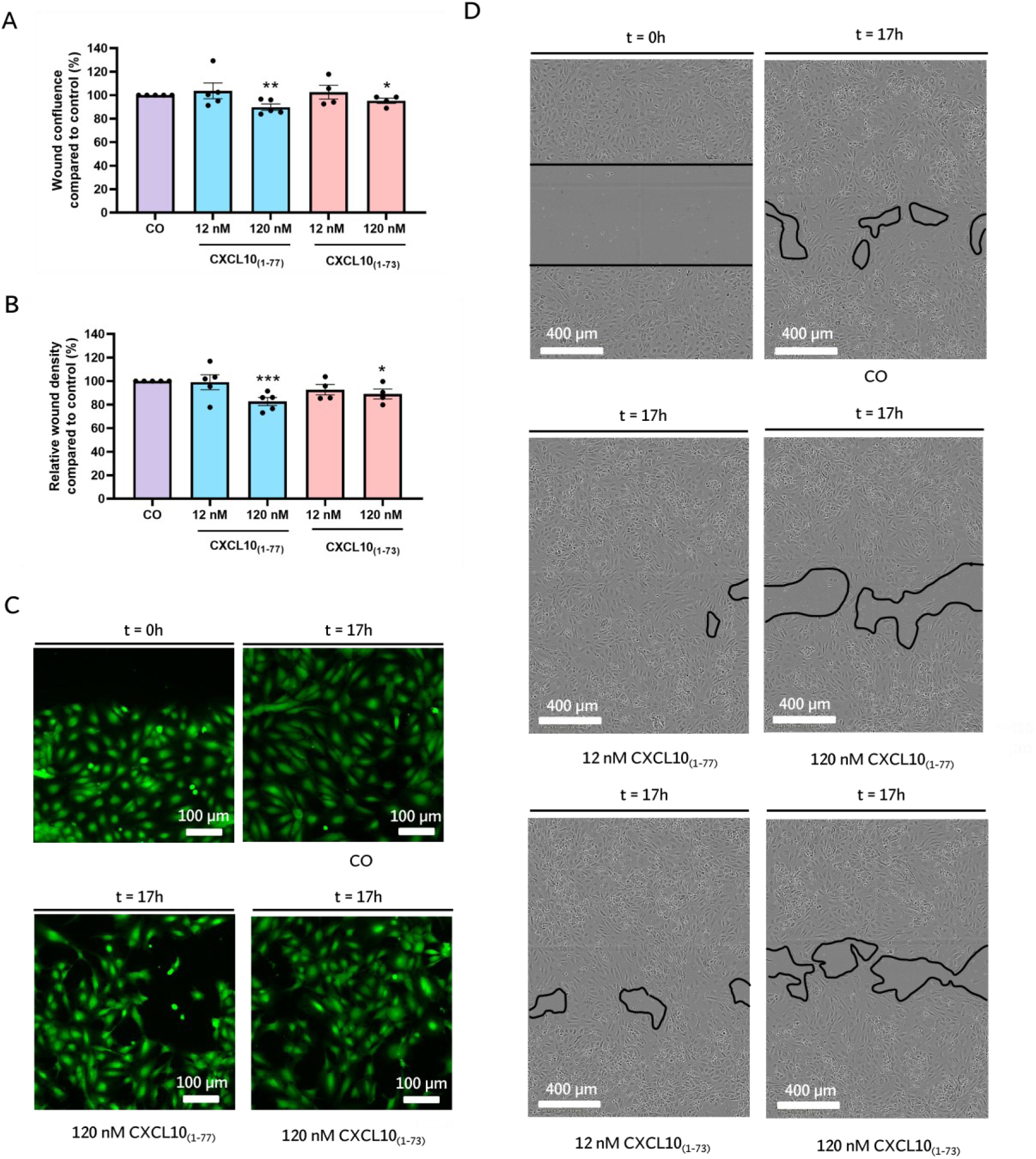
Equivalent inhibition of spontaneous HMVEC migration and invasion by intact CXCL10_(1-77)_ or C-terminally truncated CXCL10_(1-73)_. After creating a scratch wound, spontaneous HMVEC migration and invasion was monitored for 17 h in EBM-2 + 1 % FCS (CO) in the presence or absence of CXCL10_(1-77)_ or CXCL10_(1-73)_ using the IncuCyte S3 Live-Cell Analysis System. The basic analyzer unit of the Incucyte S3 2017 Software was used to calculate wound confluence and relative wound density at each timepoint. Percentages of (**A**) relative wound density and (**B**) wound confluence compared to medium-treated cells were represented in bar plots. The data are displayed as mean (± SEM) of 4 to 7 independent experiments. Unpaired t-test was used to compare differences in relative wound density and wound confluence compared to EBM-2 + 1% FCS treated cells (CO) (* p < 0.05, ** p < 0.01, *** p < 0.001). (**C**) Representative images of the wound borders using immunofluorescence microscopy after calcein staining of HMVEC stimulated with EBM-2 + 1% FCS (CO), CXCL10_(1-77)_ or CXCL10_(1-73)_ at 120 nM. Scale bar = 100 µm. (**D**) Representative images of the full wound area using IncuCyte time-lapsed microscopy pictures of HMVEC stimulated with EBM-2 + 1% FCS (CO), CXCL10_(1-77)_ or CXCL10_(1-73)_ at 12 nM and 120 nM. Scale bar = 400 µm.

Fourth, we examined the ability of CXCL10 proteoforms to blunt the FGF-2-induced ERK signal transduction pathway. CXCL10_(1-73)_ and CXCL10_(1-77)_ significantly diminished FGF-2-induced ERK phosphorylation at 120 nM with no significant differences between the two proteoforms (**Fig. 8A**). In addition, we evaluated the effects of both CXCL10 proteoforms in the *in vitro* spheroid sprouting assay, which is a solid *in vitro* angiogenesis model that is proximal to the *in vivo* situation and enables to assess angiogenesis in a 3-dimensional (3D) environment [64,65]. Pronounced sprouting of collagen-embedded HMVEC spheroids was observed after treatment with 10 ng/ml FGF-2 for 16 h (**Fig. 8B-D**). CXCL10_(1-77)_ and CXCL10_(1-73)_ at concentrations of 120 nM efficiently diminished the FGF-2-induced number of sprouts that outgrew and reduced the cumulative sprout length of spheroids, whereas both CXCL10 proteoforms were not effective at 12 nM (**Fig. 8B-D**). In summary, these *in vitro* findings demonstrate that CXCL10_(1-77)_ and CXCL10_(1-73)_ have a comparable potency to suppress spontaneous and growth factor-induced angiogenic actions including endothelial cell migration, proliferation, wound healing, signal transduction and spheroid sprouting.

**FIGURE 8.**
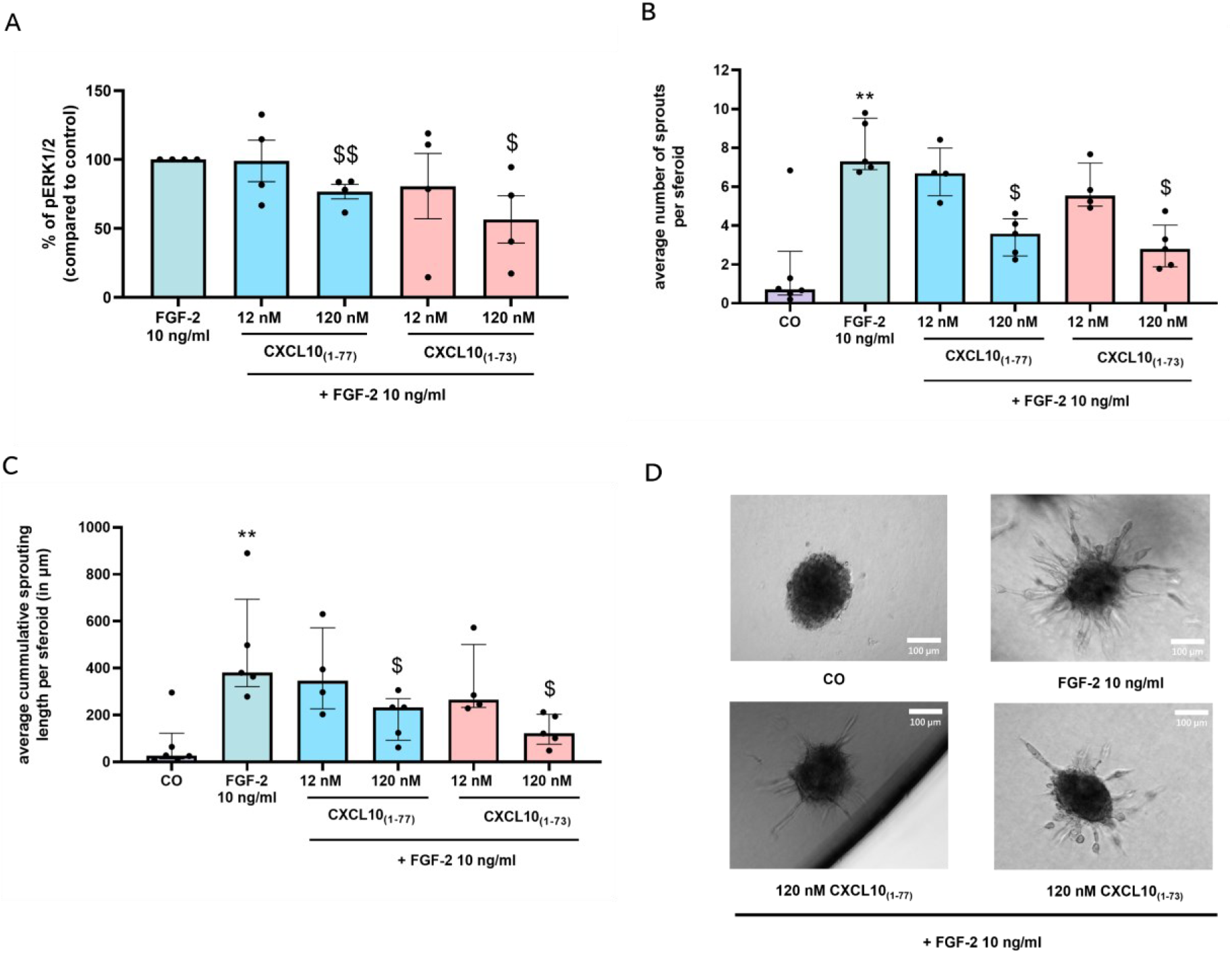
Equipotent inhibition of FGF-2-induced pERK1/2 signaling and spheroid sprouting by intact CXCL10_(1-77)_ or C-terminally truncated CXCL10_(1-73)_. (**A**) Phosphorylation of ERK1/2 was evaluated after 5 min stimulation of HMVEC with FGF-2 (10 ng/ml) in the absence or presence of CXCL10_(1-77)_ or CXCL10_(1-73)_ at 12 or 120 nM. The data are displayed as mean (± SEM) of 4 independent experiments. Unpaired t-test was performed ($ p < 0.05, $$ p < 0.01 for comparison to FGF-2). Sprouting of collagen-embedded HMVEC spheroids was assessed upon stimulation with control medium EBM-2 + 3% FCS (CO), 10 ng/ml FGF-2 alone or with CXCL10_(1-77)_ or CXCL10_(1-73)_ at the indicated doses after 16 h incubation at 37°C and 5% CO_2_. (**B**) Average number of sprouts per spheroid and (**C**) average cumulative sprouting length per spheroid (in µm) were determined with Fiji Software. The data are displayed as median (± IQR) of 4 to 5 independent experiments. Mann-Whitney U test was performed (** p < 0.01 for comparison to control, $ p < 0.05 for comparison to FGF-2). (**D**) Representative images of spheroids that were untreated (incubated with control medium EBM-2 + 3% FCS; CO), incubated with 10 ng/ml FGF-2 in the presence or absence of 120 nM of CXCL10_(1-77)_ or CXCL10_(1-73)_ are displayed. Scale bar = 100 µm.

### CXCL10_(1-73)_ induces less *in vivo* migration of CXCR3^+^ lymphocytes compared to intact CXCL10_(1-77)_

Chemokine injection into the peritoneal cavity of NMRI mice followed by peritoneal lavage was used as an experimental model to examine the *in vivo* ability of CXCL10_(1-73)_ to attract leukocytes. Mice received the competitive CD26 inhibitor sitagliptin via drinking water for 72 h prior to chemokine injection (**Fig. 9A**) to avoid CD26-mediated cleavage [35,66]. Sitagliptin thereby preserves the integrity of the N-terminus of CXCL10 and its ability to induce lymphocyte attraction after intraperitoneal (IP) injection in mice [35]. This allows clear evaluation of the effects of the C-terminal truncation. Mice had an average intake of 10 mg/day of sitagliptin via drinking water (**Fig. S7**). Soluble CD26 enzymatic activity in the peritoneal lavage fluid of sitagliptin-treated mice was significantly diminished compared to lavage fluids of untreated mice (**Fig. 9B**), confirming CD26 inhibition at the site of chemokine injection. Furthermore, immunophenotyping of peritoneal leukocytes was performed via flow cytometry (**Fig. S8**). IP injection of CXCL10_(1-77)_, but not CXCL10_(1-73)_, significantly augmented recruitment of T cells and, in particular, activated CXCR3^+^ T cells compared to vehicle-treated mice (**Fig. 9C-D**). In addition, trends towards increased ingress of CD4^+^ T cells, NKT cells, and B cells, and their activated CXCR3^+^ subsets were found for CXCL10_(1-77)_-treated mice, but not for littermates receiving CXCL10_(1-73)_ (**Fig. S9A-F**). However, lymphocyte trafficking *in vivo* can also be affected by changes in vascular permeability and lymphocyte adhesion molecules. Therefore, we assessed whether CXCL10 proteoforms influenced vascular permeability of confluent monolayers of HMVEC (**Fig. 10A**). Confluence of the monolayers on the transwell inserts was confirmed through confocal microscopy (**Fig. S10A**). Both CXCL10 proteoforms at 360 nM did not affect VEGF-induced vascular permeability (**Fig. 10A**). We also examined whether CXCL10 proteoforms altered the presence of lymphocyte adhesion molecules, tight junctions, and adherence junctions of HMVEC. PECAM-1/CD31 was significantly decreased by combined treatment with 100 ng/ml TNF-α and 100 ng/ml IFN-γ (**Fig. 10B**) as previously reported [67]. CXCL10_(1-77)_ and CXCL10_(1-73)_ did not affect the expression of PECAM-1. Combined treatment with 100 ng/ml TNF-α and 100 ng/ml IFN-γ significantly augmented the expression of lymphocyte adhesion molecules including intracellular adhesion molecule 1 (ICAM-1) and vascular cell adhesion molecule 1 (VCAM-1) (**Fig. 10C-F**). Again, both CXCL10 proteoforms did not affect expression of ICAM-1 (**Fig. 10C-D**) and VCAM-1 (**Fig. 10E-F**) upon 48 hours incubation. Furthermore, the two CXCL10 proteoforms did not affect the expression of adherence junction vascular endothelial [VE]-cadherin (**Fig. S10B-C**) nor tight junction zona occludens 1 (ZO-1) (**Fig. S10D-E**). Hence, these findings provide further evidence that CXCL10_(1-73)_ is less potent in inducing T lymphocyte chemotaxis *in vivo* compared to CXCL10_(1-77)_ by a presumable direct effect on CXCR3^+^ T lymphocytes.

**FIGURE 9.**
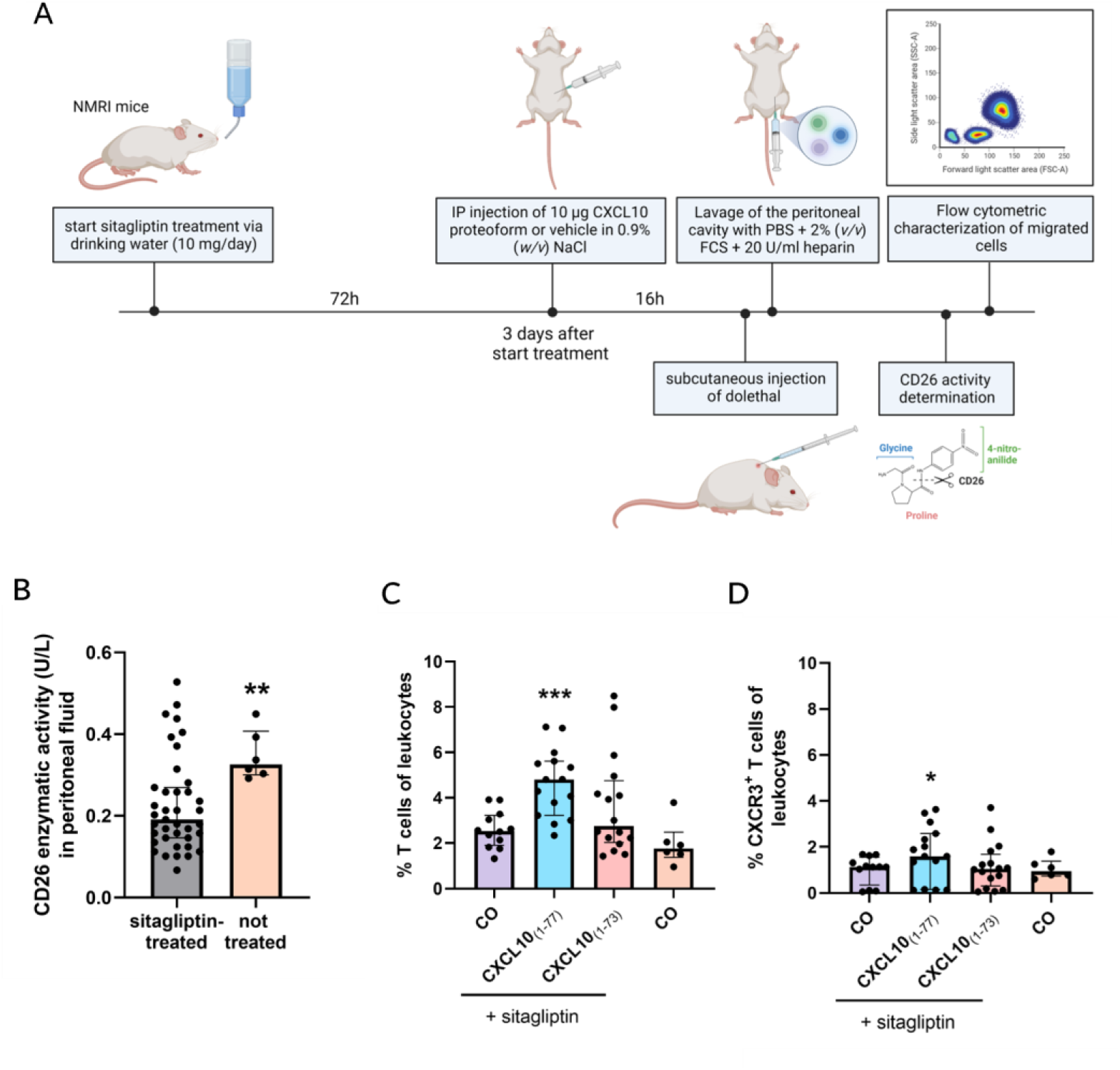
C-terminally truncated CXCL10_(1-73)_ evokes less *in vivo* migration of CXCR3^+^ T lymphocytes upon intraperitoneal injection compared to intact CXCL10_(1-77)_. (**A**) Schematic representation of the experimental set-up. Female NMRI mice received sitagliptin via drinking water for 72 h (10 mg/day) and were intraperitoneally injected with vehicle (CO), 10 µg CXCL10_(1-77)_ or 10 µg CXCL10_(1-73)_ dissolved in 0.9% (*w/v*) NaCl 16 h prior to lavage of the peritoneal cavity. Migrated cells were analyzed through flow cytometry. Inhibition of soluble CD26 (sCD26) activity in the peritoneal lavage fluids was verified in a CD26 activity assay. (**B**) sCD26 enzymatic activity (U/l) in peritoneal lavage fluids of mice treated with sitagliptin and untreated mice. (**C**) Proportions of T cells (gated as CD3^+^ NK1.1^−^) and (**D**) of activated CXCR3^+^ T cells (gated as CD3^+^ NK1.1^−^ CXCR3^+^) relative to total numbers of leukocytes (CD45^+^ cells) were determined. Each symbol represents an individual mouse (n ≥ 6 per group). Four independent experiments were performed. Horizontal lines and error bars mark the median number of cells with interquartile range. Statistical analysis was performed using a Mann-Whitney U test (* p < 0.05, ** p < 0.01, *** p < 0.001).

**FIGURE 10.**
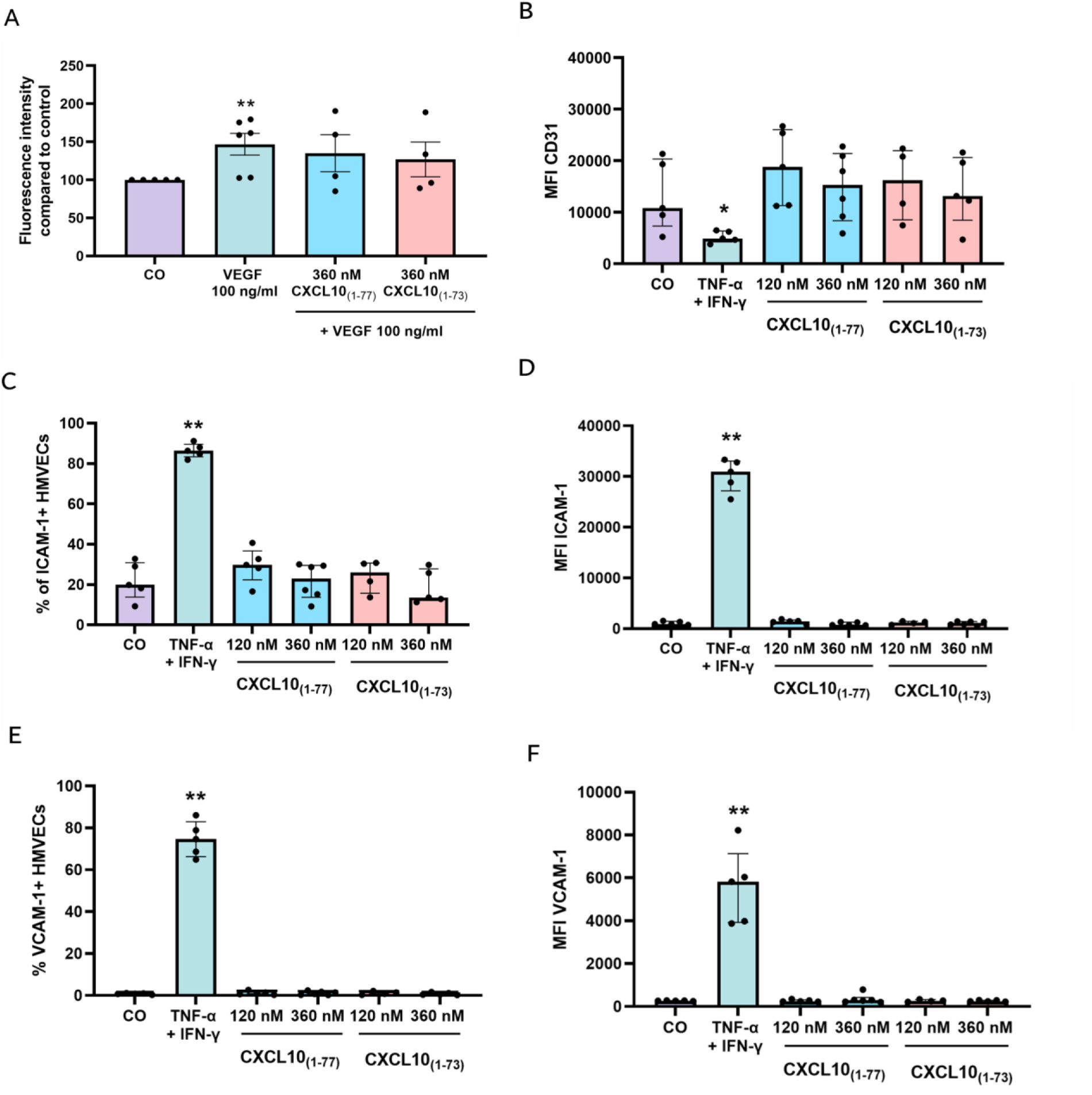
Permeability and lymphocyte adhesion molecule expression of endothelial cells is not increased by C-terminally truncated CXCL10_(1-73)_ or intact CXCL10_(1-77)_. (**A**) Endothelial monolayer permeability was assessed after stimulation with control medium EBM-2 without phenol red + 1% FCS (CO), or stimulated with VEGF (100 ng/ml) alone or VEGF (100 ng/ml) combined with 360 nM CXCL10_(1-77)_ or CXCL10_(1-73)_. Data are displayed as median (± IQR) of 4 to 6 independent experiments. Mann-Whitney U test was performed (* p < 0.05, ** p < 0.01 for comparison to control). Expression and/or MFI of (**B**) PECAM-1/CD31, (**C-D**) ICAM-1/CD54, and (**E-F**) VCAM-1/CD106 on HMVEC (gated as CD31^+^ cells) was evaluated through flow cytometry upon stimulation for 48 h at 37°C and 5% CO_2_ with control medium EBM-2 + 3% FCS (CO), 100 ng/ml TNF-α and 100 ng/ml IFN-γ, or CXCL10_(1-77)_ or CXCL10_(1-73)_ at the indicated doses. The data are displayed as median (± IQR) of 4 to 6 independent experiments. Mann-Whitney U test was performed (* p < 0.05, ** p < 0.01 for comparison to control).

## DISCUSSION

In the present study, we characterized the effects of a synthetic CXCL10 proteoform corresponding to natural C-terminally truncated CXCL10_(1-73)_ that was previously identified in human cell culture supernatant [27,29,30]. We first developed a strategy for Fmoc-based SPPS of CXCL10_(1-73)_ to ensure the availability of sufficient amounts of pure proteoform. CXCL10_(1-77)_ was previously generated through SPPS based on *tertiary* butyloxycarbonyl (*t*Boc) chemistry [58,68]. However, major drawbacks of Boc chemistry include the use of corrosive trifluoroacetic acid (TFA) in the synthesizer for removal of N-terminal Boc protection groups and the hazardous hydrofluoric acid (HF) for peptide cleavage from the solid phase support [69]. These obstacles were surmounted via Fmoc chemistry [69], whereby Fmoc protection groups are removed under moderate basic conditions and cleavage of peptides from the resin is performed via TFA. However, the moderate hydrophobic nature of CXCL10_(1-73)_ significantly hampered correct Fmoc-based SPPS, resulting in a poor yield with highly abundant contaminants consisting of incompletely synthesized peptides. Indeed, proteins comprising of a high number of amino acids possessing hydrophobic side chains are termed “difficult peptides”, given their profound challenges and complications in terms of their synthesis and purification [70,71]. These proteins tend to form inter- and intra-molecular β-sheet interactions, resulting in on-resin aggregation during peptide synthesis and consequently synthesis failure. To overcome this major obstacle, the concomitant use of pseudoproline dipeptides and a hydrophilic polyethylene glycol (PEG) resin was explored. This strategy was previously shown to substantially increase the synthesis yield of the human chemokine RANTES/CCL5_(1-68)_ [72]. In addition, we used a high quality coupling system, i.e., HCTU and NMM [55–57], as described in a recently established methodology for Fmoc-based SPPS of mCXCL10_(1-77)_ [57]. The latter authors also utilized a pseudoproline dipeptide at Ala^43^-Thr^44^, in addition to other modifications compared to the ones described in this study, as challenging protein regions in terms of SPPS differed between human CXCL10_(1-73)_ and mCXCL10_(1-77)_. For successful Fmoc synthesis of human CXCL10_(1-73)_, incorporation of an Fmoc-Ile-Pro dimer at position Ile_30_-Pro_31_ and three additional pseudoprolines (at positions Arg^5^-Thr^6^, Ile^12^-Ser^13^, and Val^68^-Ser^69^) was essential. The established SPPS approach for human CXCL10_(1-73)_ can be easily extrapolated towards other CXCL10 proteoforms (e.g., CXCL10_(3-73)_, CXCL10_(4-73)_, CXCL10_(5-73)_, CXCL10_(6-73)_, CXCL10_(1-77)_, CXCL10_(3-77)_, CXCL10_(4-77)_, CXCL10_(5-77)_, and CXCL10_(6-77)_). N-terminal truncations and an intact C-terminus can be incorporated through earlier termination of the synthesis and usage of a 2-chlorotrityl resin to prevent diketopiperazine formation associated with a C-terminal Pro [57], respectively.

A former research effort aiming to study native C-terminally truncated CXCL10_(1-73)_ was made by Hensbergen *et al.* [29]. They opted for the use of recombinant CXCL10_(1-73)_ with an additional N-terminal methionine (Met-CXCL10_(1-73)_) [29], which is an artefact due to the expression of the chemokine in bacteria. Met-CXCL10_(1-73)_ was generated via furin- and carboxypeptidase B-mediated C-terminal cleavage of recombinant Met-CXCL10_(1-77)_ [29]. Met-CXCL10_(1-73)_ retained equal potency to Met-CXCL10_(1-77)_ to induce chemotaxis of PHA- and IL-2-stimulated primary human T cells, Gα and intracellular calcium signaling in CXCR3-transfected CHO cells, and inverse agonism on the human herpes virus 8 (HHV-8)-associated ORF74 receptor [29]. In contrast, we observed significantly attenuated intracellular calcium signaling, ERK and PKB/Akt phosphorylation evoked by synthetic CXCL10_(1-73)_, indicating that second messenger signaling downstream of CXCR3A through Gαq and Gβγ is severely affected by the C-terminal truncation. These contrasting findings may be explained by the fact that CXCL10-mediated calcium mobilization and chemotaxis is known to be strongly impaired by the presence of an additional N-terminal Met in the primary sequence of CXCL10 [34]. Hence, Met-CXCL10_(1-73)_ may be an inadequate substitute for natural human CXCL10_(1-73)_. In addition, our findings accorded with data of Antonia *et al.* demonstrating that a CXCL10 proteoform lacking the α-helical and coiled C-terminal residues displayed significantly impaired chemotaxis of CXCR3^+^ Jurkat T cells *in vitro* [44].

In terms of the C-terminal residues of mCXCL10, a C-terminally mutated mCXCL10 containing K71E, R72Q, K74Q, and R75E exhibited reduced heparin and mouse CXCR3 binding affinity, diminished intracellular calcium mobilization, decreased chemotaxis of 300-19/mCXCR3 transfected cells, and an impaired ability to induce mCXCR3 internalization [47]. Mutant K71E/R72Q/K74Q/R75E mCXCL10 with an additional R22A mutation (C-tR22A) displayed even more pronounced impairment of the aforementioned functional features [47]. Furthermore, this C-tR22A mCXCL10 mutant failed to execute hallmark properties of native mCXCL10_(1-77)_, including inhibition of proliferation of human umbilical vein endothelial cells (HUVEC) [48], cell surface binding of dengue virus to mouse hepatoma cells [49] and chemotactic migration of primary lung fibroblasts treated with bronchoalveolar lavage fluid (BALF) of bleomycin-treated mice [50]. These features were attributed to the attenuated GAG binding ability of C-tR22A mCXCL10 [47]. Mature secreted human CXCL10_(1-77)_ has 70.1% amino acid identity (54/77 amino acids) with mature secreted mCXCL10_(1-77)_ and their C-terminal α-helical and coiled residues Pro^56^-Pro^77^ show profound conservation (16/22 amino acids; 72.7%; **Fig. 11A**). Hence, our findings that the loss of the two C-terminally located basic amino acids (Lys^74^ and Arg^75^) in human CXCL10_(1-73)_ diminishes the affinity for GAGs is substantiated at multiple levels. Firstly, the evolutionary conserved positively charged Lys^74^ and Arg^75^ in human CXCL10 (although not being part of a paradigmatic GAG-binding chemokine motif BBXB or BBBXXBX [73]) may serve as GAG binding residues, as shown for mCXCL10 [47]. Secondly, Lys^74^ and Arg^75^ of the C-terminus are located in proximity to the Arg^22^ of the N/20s loop and 40s loop in the structure model of CXCL10_(1-77)_, similar to mouse CXCL10 [47]. Therefore, the C-terminal amino acids would be positioned adjacent to the platelet factor 4 (PF4/CXCL4)-based predicted GAG binding residues of CXCL10_(1-77)_ (Arg^22^, Lys^46^, Lys^47^, Lys^48^, Lys^62^, Lys^66^ [46]) as shown in Fig. 11B. As such, these C-terminal residues may constitute direct GAG binding or indirectly affect the other amino acids involved in GAG binding within CXCL10 due to their vicinity. In addition to its reduced affinity for GAGs, we observed that CXCL10_(1-73)_ displayed attenuated CXCR3A signaling and diminished chemotactic potency for T lymphocytes. Booth *et al*. postulated that interaction of CXCL10 with CXCR3A would involve two demarcated hydrophobic clefts formed by the N loop and the 40s loop, in addition to the N-terminus and the 30s loop (**Fig. 11A**) [45]. More specifically, Val^7^, Arg^8^, Gln^17^, Val^19^, and Arg^38^ would be responsible for CXCR3A binding based on the NMR structure (**Fig. 11C**) [45]. Additional partially overlapping CXCR3 binding regions were identified by others, including Asn^20^-Cys^36^ [46] and Arg^8^-Pro^21^ and Glu^40^-Gly^49^ [74] (**Fig. 11C**). Thus, Arg^22^ (20s loop), Lys^46^, Lys^47^ and Lys^48^ (40s loop) are likely involved in both GAG and receptor binding of CXCL10. Moreover, the initial high affinity binding of CXCL10 to CXCR3A is dependent on interaction with negatively charged sulfated Tyr^27^ and Tyr^29^ and N-glycosylated Asn^22^ and Asn^32^ in the N-terminal region of CXCR3A [75–77]. Hence, we hypothesize that the spatial vicinity of the positively charged C-terminal residues Lys^74^ and Arg^75^ and the N/20s-loop (in particular Arg^22^ implied in CXCR3A binding [46]) may form a Coulomb-assisted interaction surface to bind sulfo-Tyr^27/29^ and N-glycosylated Asn^22/32^ and thereby enable docking to CXCR3A. Presumably, the absence of C-terminal Lys^74^ and Arg^75^ would result in hampered or delayed docking to CXCR3A. Consistent with this hypothesis, we observed a delay in the initiation of calcium responses to CXCL10_(1-73)_ (**Fig. 4B**), which may be explained by delayed docking. Thus, the C-terminal residues of human CXCL10 may act analogous to mCXCL10 where Lys^71^-Arg^75^ are involved in CXCR3A activation resulting in downstream calcium signaling and chemotaxis (*vide supra*) [47]. Also, similar to mCXCL10 is the emerging notion that binding sites for GAG and CXCR3 are partially overlapping in human CXCL10.

**FIGURE 11.**
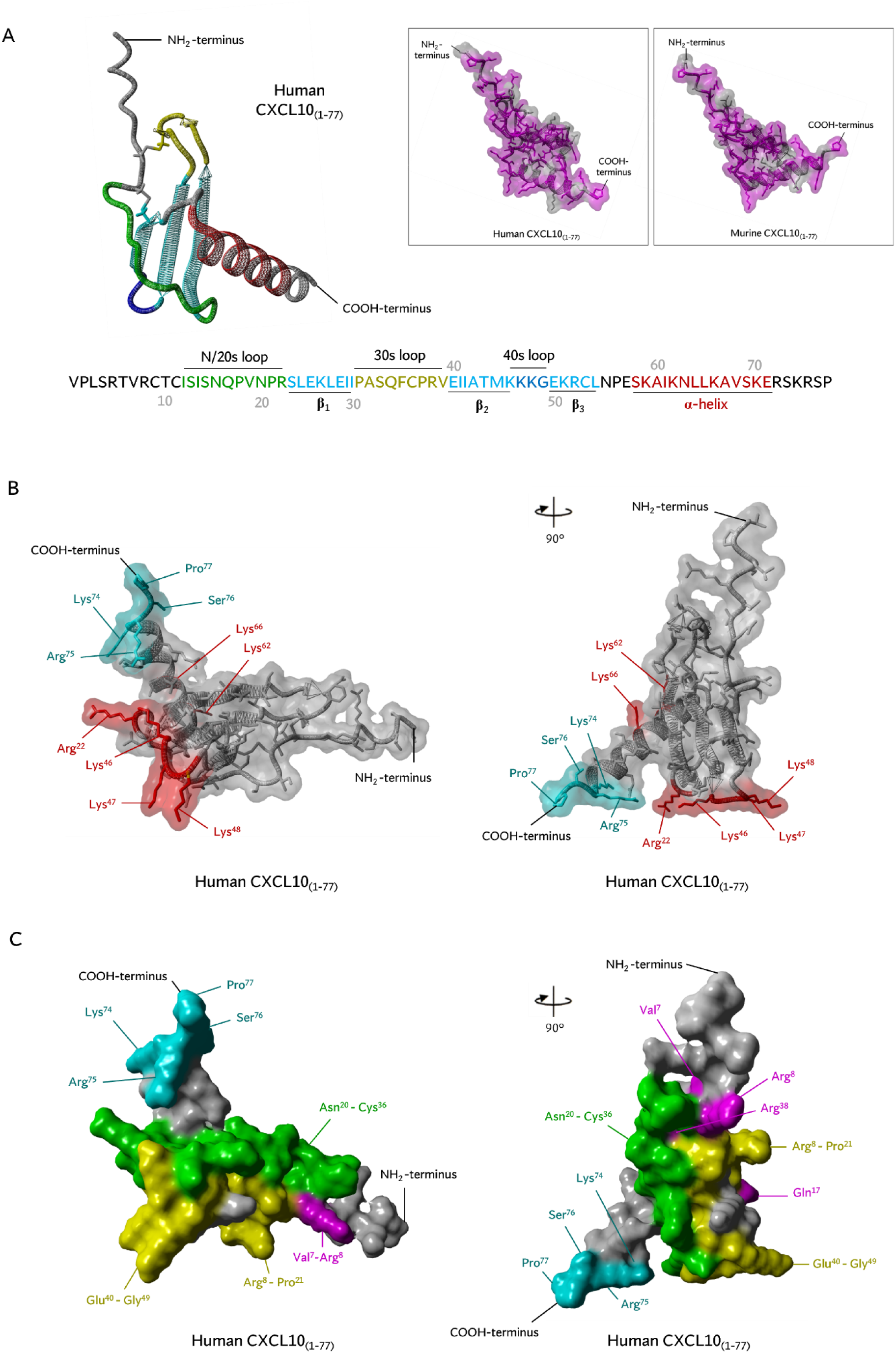
Structure models of human CXCL10_(1-77)_. (**A**) Structure models of human CXCL10_(1-77)_ of AlfaFold DB (right; AF-P02778-F1 without signal sequence). This model is based on the crystal structure of CXCL10 in hexagonal (H) form [PDB accession code 1O80], crystal structure of CXCL10 in monoclinic (M) form [PDB accession code 1O7Y], crystal structure of CXCL10 in tetragonal (T) form [PDB accession code 1O7Z], and NMR spectroscopy-determined CXCL10 [PDB accession code 1LV9] [45,46] whereby unobserved Ser^76^ and Pro^77^ were predicted through AlfaFold DB. Secondary structures of human CXCL10_(1-77)_ are displayed. CXCL10 has an N/20s loop (green), three antiparallel β-sheets (cyan), 30s loop (yellow), 40s loop (blue), and an α-helix (red). The inset shows human CXCL10_(1-77)_ (AF-P02778-F1) and mouse CXCL10_(1-77)_ of AlfaFold DB (left; AF-Q3UK71-F1 without signal sequence). Conserved residues in mouse CXCL10_(1-77)_ and human CXCL10_(1-77)_ (magenta) and residues that are not conserved (grey) are displayed. (**B**) The structural model of human CXCL10_(1-77)_ of AlfaFold DB (AF-P02778-F1) is shown from two different perspectives with a transparent surface to visualize amino acid side chains. Four C-terminally residues that are shedded in CXCL10_(1-73)_ (cyan) are located in close proximity to predicted potential GAG-binding residues Arg^22^, Lys^46^, Lys^47^, Lys^48^, Lys^62^, and Lys^66^ (red) [46]. (**C**) The structural model of human CXCL10_(1-77)_ of AlfaFold DB (AF-P02778-F1) is displayed from two different perspectives with a non-transparent surface to visualize the receptor interaction surface. Potential CXCR3 binding residues of CXCL10 are indicated in different colors as previously shown by Swaminathan *et al.* [46]: residues of human CXCL10_(1-77)_ found to be perturbed in 2D ^15^N-^1^H HSQC NMR spectra by the addition of an N-terminal CXCR3 peptide CXCR3_(22-42)_ [45] (Val^7^, Arg^8^, Gln^17^, Val^19^, and Arg^38^; magenta), residues bound by a CXCR3-blocking anti-CXCL10 monoclonal antibodies preventing chemotaxis and calcium mobilization [46] (Asn^20^-Cys^36^; green), and residues aligned to the CXCL8 binding region to CXCR1 (Arg^8^-Pro^21^ and Glu^40^-Gly^49^; yellow) [74]. Four C-terminally residues that are shedded in CXCL10_(1-73)_ (cyan) are positioned in vicinity of several potential CXCR3 binding residues.

Analogous to CD26-mediated N-terminally truncated CXCL10_(3-77)_ [34], we observed that the C-terminal truncation did not significantly modulate anti-angiogenic properties of CXCL10. Hence, proteolysis of CXCL10 in inflamed tissue would still favor angiostasis, thereby limiting additional leukocyte ingress in the respective epicenter of inflammation by preventing formation of new blood vessels. Whether the retained angiostatic effect of CXCL10_(1-73)_ is mediated through the proposed angiostasis-mediating CXCR3B [78], through the unidentified receptor for CXCL10 on endothelial cells [79], through differential PTMs of CXCR3 on endothelial cells compared to leukocytes, or through a biased cell type-dependent CXCR3A or GAG signaling pathway [25,48] (that is not affected by these truncations), remains elusive. Intriguingly, Gao *et al*. reported that the N-terminal domain of CXCR3B lacked sulfo-Tyr residues [76], whereas others have reported that the extended N-terminus of CXCR3B has two additional sites (Tyr^6^ and Tyr^40^) that may be prone to sulfation [80,81]. Moreover, lymphocytes exhibit less heparan sulfate proteoglycans on their surface compared to other leukocytes [82], which may emphasize the importance of sulfation sites on CXCR3A to sequester chemokines [75]. In contrast, endothelial cells have relatively more GAGs expressed on their surface, again pointing towards potentially differential signaling of CXCL10 in HMVECs compared to T cells. Alternatively, CXCL10 may also interfere with binding of growth factors to their receptors by directly binding to the growth factor or impairing the growth factor’s ability to dimerize, as shown for CXCL4 [83]. Given that the CXCL10-derived peptide CXCL10_(56-77)_ exerted similar anti-angiogenic effects as full-length CXCL10_(1-77)_ [43], our findings may also imply that the C-terminal α-helical amino-acids (Ser^58^-Glu^71^; **Fig. 11A**) —rather than the endmost C-terminal coiled residues— are crucial for actions leading to downstream signaling resulting in angiostasis.

In general, processing of CXCL10_(1-77)_ into CXCL10_(1-73)_ with reduced GAG-binding affinity and decreased receptor signaling potency attenuates CXCL10-mediated leukocyte infiltration whilst facilitating angiostasis. However, GAG binding affinity of CXCL10_(1-73)_ was not equivalently attenuated for different GAGs (HS > CS-A > heparin), which may have physiological implications for the actions of CXCL10. In tissues where GAGs are expressed for which CXCL10_(1-73)_ has less affinity and both CXCL10 forms occur, the net-effect of reduced CXCL10_(1-73)_ immobilization on GAGs may enhance the CXCL10_(1-77)_-mediated chemotactic effect as (a) more GAG binding sites are available for functional CXCL10_(1-77)_ and (b) less CXCL10_(1-73)_ (which is less potent) is able to bind CXCR3A^+^ leukocytes. For example, proteolysis of CXCL10_(1-77)_ into CXCL10_(1-73)_ probably occurs in inflamed joints as the C-terminally truncating enzymes furin, carboxypeptidase B, MMPs, and cathepsins are expressed in the synovium [84–90]. Distinct distribution of GAGs in the synovium compared to the blood circulation may thus impact localized functioning of CXCL10.

In conclusion, our study reveals that the C-terminal residues Lys^74^-Pro^77^ of CXCL10 are important for GAG binding, CXCR3A signaling, T lymphocyte chemotaxis, but dispensable for angiostasis. In addition, the optimized SPPS approach to generate high quality synthetic CXCL10 paves the way towards research on other naturally occurring CXCL10 proteoforms. Given the validated role of CXCL10 in viral infection [37–41,91,92], tumor immunology [36,93,94] and autoimmune arthritis [95,96], the balance between CXCL10 and its processing enzymes (e.g. CD13, CD26, furin, carboxypeptidase B, MMPs, and cathepsins) in inflamed tissues is pivotal for fine-tuning the effects of CXCL10 in (patho)physiological settings.

## MATERIALS AND METHODS

A Supplemental Experimental Procedure section is provided in the Supplemental Information.

### Cell cultures and reagents

Chinese hamster ovary (CHO) cells transfected with CXCR3A were cultured in Ham’s F-12 growth medium (Gibco; Thermo Fisher Scientific, Waltham, MA, USA) supplemented with 10% (*v/v*) heat-inactived fetal calf serum (FCS, Sigma-Aldrich, Saint Louis, MO, USA), 400 µg/ml G418 (Carl Roth, Karlsruhe, Germany), 1 mM sodium pyruvate (Gibco) and 0.12% (*v/v*) sodium bicarbonate (Gibco) [34]. Human microvascular endothelial cells (HMVEC; Cell Systems, Kirkland, WA, USA) were cultured in endothelial cell basal medium-2 (EBM-2: Lonza, Basel, Switzerland) supplemented with the EGM-2 MV SingleQuots kit (Lonza). For culturing of primary lymphocytes, PBMC were purified from buffy coats of healthy volunteers (Red Cross, Mechelen, Belgium) through gradient centrifugation, as previously described [97]. T lymphoblasts were generated through culturing mononuclear cells in 2 µg/mL PHA (Sigma-Aldrich) and 50 U/ml interleukin (IL)-2 (PeproTech, Rocky Hill, NJ, USA), as formerly described [34].

### Chemical synthesis and purification of C-terminally truncated human CXCL10_(1-73)_

CXCL10_(1-73)_ was chemically synthesized based on *N*-(9-fluorenyl)methoxycarbonyl (Fmoc) chemistry using an Activo-P11 automated peptide synthesizer (Activotec, Cambridge, UK). A hydrophilic resin and specialized amino acid building blocks were used to ensure a successful SPPS (**Figure 1**).

### Surface plasmon resonance

Real-time binding kinetics of CXCL10 proteoforms with different GAGs (heparin, HS, and CS-A) were examined through SPR on a BIAcore T200 instrument (Cytiva, Uppsala, Sweden) in a similar experimental set-up as previously described [98].

### Signal transduction assays

The potency of CXCL10_(1-77)_ and CXCL10_(1-73)_ to induce an increase of the intracellular calcium concentration was evaluated on CXCR3A-transfected CHO cells, as previously described [34,99]. To determine phosphorylation of ERK1/2 and Akt upon chemokine treatment, 0.4 × 10^6^ CXCR3A-transfected CHO cells/ml or 60 000 HMVEC/ml (2ml/well) were seeded in flat bottom 6-well plates (2.0 ml/well; TPP, Sigma-Aldrich) in supplemented Ham’s F-12 growth medium + 10% (*v/v*) FCS or EBM-2 cell culture medium, respectively. Upon overnight starvation in serum-free medium, cells were incubated with serum-free medium containing 0.5% (*w/v*) bovine serum albumin (BSA; endotoxin free, Sigma-Aldrich) for 15 min at 37°C. Subsequently, cells were stimulated at 37°C with CXCL10_(1-73)_ or CXCL10_(1-77)_ for 5 min (for CHO cells) or 15 min followed by 5 min stimulation with FGF-2 (for HMVEC). Signal transduction was terminated and pERK1/2 and pAkt was determined in the supernatant of cell lysates, as previously described [27,99].

### Multiscreen chemotaxis assay of primary T lymphocytes

For the multiscreen chemotaxis assay (Millipore Corporation, Billerica, MA, USA), 96-well filter plates (5 µm pore-size) were either not pre-coated or pre-coated with bovine plasma FN (Gibco), human FN (BD Biosciences, San Jose, California, USA) or human type I collagen (Sigma-Aldrich) overnight. Primary T lymphocytes stimulated with PHA and IL-2 (2 × 10^5^, 100 µl/well) were resuspended in HBSS buffer (Gibco) containing 0.1% (*w/v*) BSA and 100 µM of the CD26 inhibitor sitagliptin (Januvia; Merck Sharpe & Dohme [MSD] Whitehouse Station, NJ, USA). T lymphocyte migration from the upper plate towards chemoattractant solution in the receiver plate was quantified via the ATP detection assay (Perkin Elmer, Waltham, MA). In parallel with the multiscreen assay, CXCR3 expression on PHA- and IL-2-activated T lymphocytes was evaluated through flow cytometry.

### CXCR3 internalization on primary T lymphocytes

Equal volumes of PHA- and IL-2 stimulated T cells (90 µl, 5.5 × 10^6^ cells/ml) were resuspended in PBS containing 100 µM sitagliptin (Januvia) and 2% FCS and stimulated with varying concentrations of CXCL10_(1-73)_ and CXCL10_(1-77)_ for 10 min at 37°C. After incubation, cells were put on ice and washed once with ice-cold flow cytometry buffer. After centrifugation at 4°C for 5 min at 300*g*, cells were resuspended in PBS. Internalization of CXCR3 was analyzed with flow cytometry in a similar manner as described to evaluate the CXCR3 expression on T lymphocytes used for multiscreen chemotaxis assays (*vide supra*). The relative surface expression of CXCR3 was calculated as a percentage relative to medium treated cells.

### xCELLigence chemotaxis assay for HMVEC

The xCELLigence® real-time cell analyzer double plate (RTCA-DP) system (ACEA Biosciences, Inc.; San Diego, CA, USA) was utilized to evaluate HMVEC migration. Briefly, 160 µl of control medium (i.e., EBM-2 medium containing 0.4% [*v/v*] FCS) with or without 30 ng/ml FGF-2 was added to the lower chamber of a cell invasion/migration (CIM) plate in the presence or absence of CXCL10_(1-77)_ or CXCL10_(1-73)_ at varying concentrations (1.2 nM, 12 nM, 120 nM, or 360 nM). Upon chamber assembly and addition of HMVEC (4 × 10^4^ cells/well) to the upper compartment, alterations in electrical impedance were measured and converted into cell indices. To compare HMVEC migration induced by CXCL10_(1-73)_ and CXCL10_(1-77)_ relative to control medium or FGF-2-treated cells at 12 h, cell indices measured upon incubation with control medium were set to 100%.

### *In vitro* toxicity assay

HMVEC were seeded at 8 × 10^3^ cells/well in MCDB131 medium + 3% (*v/v*) FCS in a black, clear bottom 96-well plate (Greiner Bio-one, Kremsmünster, Austria) coated with 0.1% (*v/v*) gelatin in PBS. Following overnight incubation (37°C, 5% CO_2_), cells were washed and incubated with control medium (*i.e*., MCDB131 supplemented with 0.4% (*v/v*) FCS) in the presence or absence of CXCL10_(1-77)_ or CXCL10_(1-73)_ for 30 h at 37°C and 5% CO_2_. Toxicity of the CXCL10 proteoforms was evaluated using the LIVE/DEAD Viability/Cytotoxicity Kit for mammalian cells (Invitrogen, Thermo Fisher Scientific) according to the manufacturer’s instructions

### Proliferation assay

HMVEC were seeded at 5 × 10^3^ cells/well in EBM-2 cell culture medium in a flat bottom 96-well plate (Greiner Bio-One). After overnight settling of the cells, cells were starved for 4 hours in EBM-2 + 1% [*v/v*] FCS. After starvation, cells were stimulated with FGF-2 (10 ng/ml) alone or in combination with CXCL10 proteoforms. The ATP lite assay (Perkin Elmer) was used according to the manufacturer’s instruction after 4 days to assess proliferation.

### Scratch wound healing assay

HMVEC were seeded at 15 × 10^3^ cells/well in EBM-2 cell culture medium in an IncuCyte ImageLock 96-well plate (Essen Bioscience; Newark, UK) coated with 0.1% (*v/v*) gelatin in PBS. Following overnight incubation (37°C, 5% CO_2_), 700 – 800 µm wide wounds were simultaneously created in the endothelial monolayers of all wells using an IncuCyte 96-well Woundmaker Tool (Essen Bioscience). Cells were washed twice in basal EBM-2 medium and incubated with control medium (*i.e*., EBM-2 + 1% [*v/v*] FCS), 1 ng/ml FGF-2 or CXCL10_(1-77)_ or CXCL10_(1-73)_ in the presence or absence of 1 ng/ml FGF-2. HMVEC were monitored for 20 h in the IncuCyte S3 Live-Cell Analysis System to determine wound confluence and relative wound density.

### Spheroid sprouting assay

Single spheroids were formed in hanging droplets, collected and distributed over a clear flat bottom 96-well plate in a methylcellulose/collagen type I suspension as previously described [98]. Spheroids were left untreated (addition of control medium, i.e. EBM-2 + 3% [*v/v*] FCS) or were incubated with 12 nM or 120 nM CXCL10_(1-77)_ or CXCL10_(1-73)_ at 37°C and 5% CO_2_ for 15 min prior to the addition of 10 ng/ml FGF-2 in EBM-2 + 3% (*v/v*) FCS. Following 17 h incubation at 37°C and 5% CO_2_, sprouting of the spheroids was evaluated with bright field imaging through a 10× objective on an inverted Axiovert 200M microscope (Carl Zeiss Microscopy GmbH, Oberkochen, Germany). The average number of sprouts per spheroid and the average cumulative sprout length per spheroid for each well was calculated using ImageJ software (NIH; Bethesda, Maryland, USA).

### *In vivo* cell migration assay

Drinking water of 8-week old Naval Medical Research Institute (NMRI) mice was supplemented with 1.7 mg/ml of CD26 inhibitor sitagliptin (Januvia) for 72 h prior to an intraperitoneal (IP) injection of 10 µg recombinant CXCL10_(1-77)_ or synthetic CXCL10_(1-73)_. Drinking volume was monitored daily. A *Limulus* amoebocyte lysate assay (Cambrex Corporation, East Rutherford, NJ, USA) showed that CXCL10_(1-73)_ and CXCL10_(1-77)_ stock solutions contained very low endotoxin levels (<0.06 pg LPS/µg of chemokine). Mice were euthanized with a subcutaneous injection of 300 µl Dolethal (pentobarbital; 200 mg/ml; Vétoquinol, Aartselaar, Belgium) 16 h after chemokine injection, and peritoneal cavities were washed with 5 ml PBS supplemented with 2% (*v/v*) FCS and 20 U/ml heparin (Leo Pharma, Amsterdam, the Netherlands). Cells were analyzed through flow cytometry.

### CD26 activity assay

After collection of cells for analysis by flow cytometry, peritoneal lavage fluids of NMRI mice were centrifuged at 300g for 10 min at 4°C and supernatant was collected and stored at −20°C. In a flat bottom 96-well plate, lavage fluids (1/2 diluted; 100 µl) were incubated with 500 µM Gly-Pro-*p*-nitroanilide substrate (Sigma-Aldrich) in 200 mM Tris-HCl buffer (pH 8.3) to determine CD26 enzymatic activity.

### Vascular permeability assay

HMVEC were seeded (10 000 cells/well) on gelatin-coated membranes with 0.4 µm pores and 6.5 mm diameter (Transwell; Corning, New York) and were grown to confluence in EBM-2 cell culture medium. After starving the cells overnight in EBM-2 + 1% [*v/v*] FCS, cells were treated with 100 ng/ml VEGF (Biolegend; San Diego, California, USA) alone or in combination with 360 nM CXCL10_(1-77)_ or CXCL10_(1-73)_ in the upper chambers for 3h. Afterwards, leakage of 1 mg/ml fluorescein isothiocyanate (FITC)-conjugated dextran (70 kDa; Sigma-Aldrich) from the top to the bottom compartment was used to calculate permeability. To check whether cell confluence was obtained, we seeded a 96-well plate in parallel at equal cell density (well surface is identical to the surface of a transwell insert) and we performed confocal microscopy on the inserts.

### Evaluation of expression of lymphocyte adhesion molecules and junctions on endothelial cells

HMVEC were seeded (140 000 cells/well) in a flat bottom 6-well plate in EBM-2 cell culture medium. Upon adherence, HMVEC were treated for 48 h at 37°C and 5% CO_2_ with EBM-2 medium + 3% FCS (CO), a combination of 100 ng/ml TNF-α (Peprotech) and 100 ng/ml IFN-γ (Peprotech) to induce lymphocyte adhesion molecules, or CXCL10_(1-77)_ or CXCL10_(1-73)_ at 120 nM or 360 nM. Thereafter, medium was removed and cells were washed in cold PBS. Cells were detached using cell scrapers (Sarstedt, Darmstadt, Germany) in 100 µl cold PBS to avoid trypsinization. Cells were transferred to FACS tubes, stained and analyzed.

### Statistical analysis

GraphPad Prism software 9.3.0 was used for data analysis. Shapiro-Wilk test was used to assess if data were normally distributed. A Mann-Whitney U test or Kruskal-Wallis test with Dunn’s multiple comparison correction was used for data that were not normally distributed (displayed as median ± IQR). Unpaired t-tests were used when the data exhibited normal distribution (displayed as mean ± SEM).

## AUTHOR CONTRIBUTION

LD performed the experiments with the help of KY, ADZ, SN, MG, NB, SB and EM. LD analyzed the data together with KY, ADZ, SN, MG, NB, DS, SS and PP. PP, SS and DS supervised the study. LD wrote the initial manuscript. All authors contributed to the study conception and design, provided their comments on different versions of the manuscript and approved the final version of the manuscript.

## ACKNOWLEDGEMENTS

Schematic representations were created with Biorender.com.

